# Mechanistic Constraints on ClpM Expression Underlie Apicoplast Genome Retention in Malaria Parasites

**DOI:** 10.64898/2026.05.20.726455

**Authors:** Alia Qasem, Shirly Kats Galay, Aseel Ghanaiem, Hari Shankar, Michal Shahar, Anat Florentin

**Author notes:** Indian Council of Medical Research, New Delhi, India (Hari Shankar). **Corresponding Author:** Anat Florentin.

## Abstract

The apicoplast of malaria parasites retains a reduced genome encoding a small set of genes with unknown functions. Among these genes is a putative ClpM chaperone, which unlike other apicoplast Clp-family members is not nuclear but plastid-encoded. In this study, we used ClpM as a model case to investigate evolutionary and molecular basis for plastid genome retention.

Phylogenetic analyses across plastid-containing eukaryotes revealed that ClpM orthologues are broadly conserved and consistently plastid-encoded in all organisms with a red alga-derived plastid, irrespective of parasitism, photosynthesis or physiology. This broad phenomenon suggested gene-specific evolutionary constraints that were subsequently tested experimentally.

To test whether clpM can be functionally expressed from the nucleus, we generated transgenic parasites carrying a nuclear ClpM copy fused to an apicoplast-targeting transit peptide. Unexpectedly, standard transgenesis resulted in transcriptional silencing, and we therefore forced transcription using integration into an endogenous essential locus. This led to robust clpM mRNA, however no detectable ClpM protein was observed. Multiple analyses ruled out apicoplast-dependent instability, ER-associated degradation, misfolding or membrane sequestration. Attempts to express clpM or other plastid-derived genes using endogenous sequences were found to be toxic, suggesting nucleotide-sequence incompatibility. In contrast, a transgene carrying a second copy of the nuclear ClpC ortholog was readily expressed. Comparative analysis of ClpM and ClpC domain architecture showed that their ATPase domains form distinct evolutionary clusters, suggesting conserved but functionally divergent roles. Subsequently, domain-swap experiments between ClpC and ClpM rescued partial expression and identified specific domains as contributors to the nuclear-expression barrier.

Together, these findings demonstrate that clpM retention in the apicoplast genome is enforced by multilayered constraints involving evolutionary conservation, nucleotide-sequence incompatibility, transcriptional block and protein-intrinsic translational barriers. This work provides experimental evidence for mechanisms that restrict organelle-to-nucleus gene transfer and contribute to organelle genome retention.

## Introduction

Malaria, one of the world’s most consequential infectious diseases, is caused by eukaryotic parasites from the genus *Plasmodium*, with one species, *P. falciparum*, accounting for the majority of severe morbidity and mortality. The global spread of drug-resistant parasites underlies the importance of identifying novel drug targets for improved therapeutic interventions^1,2^.

A central reason that *Plasmodium* parasites are vulnerable to selective antibiotics is the presence of the apicoplast: a relict, non-photosynthetic plastid that houses prokaryote-like biosynthetic pathways^3,4^. The apicoplast has retained bacterial-type gene expression, consistent with the fact that antibiotics targeting prokaryotic translation compromise the organelle and kill parasites with “delayed-death” kinetics^5,6^. In asexual blood stages of *P. falciparum*, the best-established essential apicoplast output is the production of isopentenyl pyrophosphate (IPP) via the MEP (non-mevalonate) pathway^7^. Chemical-rescue experiments showed that parasites survive loss of the apicoplast when supplied with exogenous IPP, underscoring isoprenoid biosynthesis as the critical blood-stage function of the organelle^8^. However, isoprenoid production is not the sole metabolic function of the apicoplast^9^; among the many apicoplast-resident metabolic pathways, some become essential during the parasite’s liver and mosquito stages (e.g. fatty acids^10^ and heme biosynthesis^11^), some are required for MEP enzymes functions (Fe–S cluster assembly^12^), and others remain active and essential even after apicoplast ablation (co-enzyme A maturation^13^). Thus, apicoplast functions cannot be reduced to IPP production, and the dependency between the *Plasmodium* cell and its ancient endosymbiont is more complex than is often being argued.

Most proteins required for apicoplast metabolism and maintenance are encoded in the nucleus, synthesized in the ER, and imported into the organelle using N-terminal targeting sequences^14,15^. Despite this streamlined transport, the apicoplast retains its own small genome^16^. The *P. falciparum* plastid genome is a ∼35 kb DNA circle with greatly reduced sequence complexity that encodes predominantly gene-expression components, including rRNAs, tRNAs, subunits of a bacterial-type RNA polymerase, multiple ribosomal proteins, and a translation elongation factor^17^. This organization highlights a core paradox of apicoplast biology: essential metabolic capacity is specified by nuclear genes, yet a minimal plastid genome has been evolutionary conserved.

Why do non-photosynthetic plastids retain genomes at all? Endosymbiont-to-organelle evolution involved extensive plastid genome reduction and gene transfer to the nucleus, but this process often stalls with a small gene set that remains organelle-encoded. Classic explanations for plastid genome retention have been proposed for plants and algae, and thus they are photosynthesis-centric; they rationalize the advantage of encoding photosystems components from the chloroplast genome for coordinated regulation coupled with organelle redox state^18,19^. These are interesting theories that remain to be experimentally-tested, however they provide no explanation for heterotrophic plastids such as those of *Plasmodium* and its close relatives, the parasites belonging to the phylum of Apicomplexa. It was previously suggested that apicomplexans retain plastid genomes because they still depend on a very small number of plastid-encoded genes, and that plastid genome loss becomes feasible when these key genes are transferred to the nucleus^20^. But what these genes are, and whether they can be nuclear-transferred is unknown.

Notably, among the very few plastid genes that do not encode transcription/translation machinery, the apicoplast genome encodes an unusual Clp-family chaperone (historically annotated as ClpC, and later defined as ClpM^21^). The Clp system provides an especially compelling framework to interrogate these constraints because it couples organelle proteostasis to developmental and metabolic demands^22^. Clp chaperones (sometimes called HSP100) form a conserved family of AAA+ ATPases that remodel protein complexes, and in many contexts cooperate with proteases to promote regulated proteolysis^23^. In *P. falciparum*, multiple Clp chaperones have been catalogued and localized to the apicoplast, including ClpB, ClpC, and a plastid-encoded ClpM/ClpC-like protein^21,22^. Together with the apicoplast ClpP/R protease core, they were suggested to form a plastid-centered protein quality control network^22,24^. Consistent with this, it was demonstrated that the apicoplast Clp system is essential for organelle biogenesis and proteostasis, with the nuclear-encoded chaperone ClpC required for complex stabilization and plastid integrity^25^. Conversely, the function of the plastid-encoded chaperone, often referred to as ClpM to distinguish it from the nuclear ClpC, is unknown. Moreover, ClpM sits at the intersection of two key problems: how apicoplast proteostasis is maintained, and why some plastid genes resist relocation to the nucleus.

In this work, we sought to experimentally investigate the evolutionary retention of the apicoplast genome in *P. falciparum*, hypothesizing that it is driven by at least one essential gene whose expression is confined to the organelle. We further propose that this cryptic gene might be the apicoplast-encoded ClpM, and therefore focused on its potential nuclear-transfer. Using a framework of nuclear transgenes under various expression systems, we demonstrate that plastid-derived genes become toxic when nuclear encoded. We found that the molecular basis preventing nuclear expression involves multilayer mechanisms, including nucleotide sequence constraints, transcriptional silencing and protein-intrinsic determinants that block translation. These findings provide experimental evidence for evolutionary forces maintaining plastid genome retention of ClpM, and reveal previously unrecognized post-transcriptional barriers governing organellar gene compartmentalization.

## Results

### Clp Chaperone is conserved and plastid-encoded in apicomplexan as well as related and ancestral clades

To understand the evolutionary context of the apicoplast-encoded Clp chaperone in *Plasmodium*, we investigated its phylogenetic conservation and genomic location across plastid-containing eukaryotes. To do that, we searched all 15,231 plastid organelle genomes available in NCBI, with *P. falciparum* ClpM (PF3D7_API03600) as query (Fig. 1A). Notably, ClpM orthologs were readily identified in a wide range of organisms with red-algae-derived plastids, while being completely absent from green plant plastids (Fig. 1B). These includes Rhodophyta, the primary red algal lineage, as well as all lineages that acquired their plastid secondarily from red algae^26^: Apicomplexa^27^, Stramenopiles^28,29^, Haptista^26^, Cryptophyta^30^, Dinoflagellata^31,32^, and Chromerids^33^ (Fig. 1B). This demonstrates that the Clp chaperone is widely conserved both in sequence and in extranuclear genomic location. Moreover, its genomic localization is not restricted by a parasite-specific function, nor constrained by photosynthesis. It is plastid-encoded in both unicellular and multicellular organisms and is unrelated to a specific ecological niche. This suggests that this genomic localization is functionally critical in rhodophyte-related plastids.

**Figure 1.**
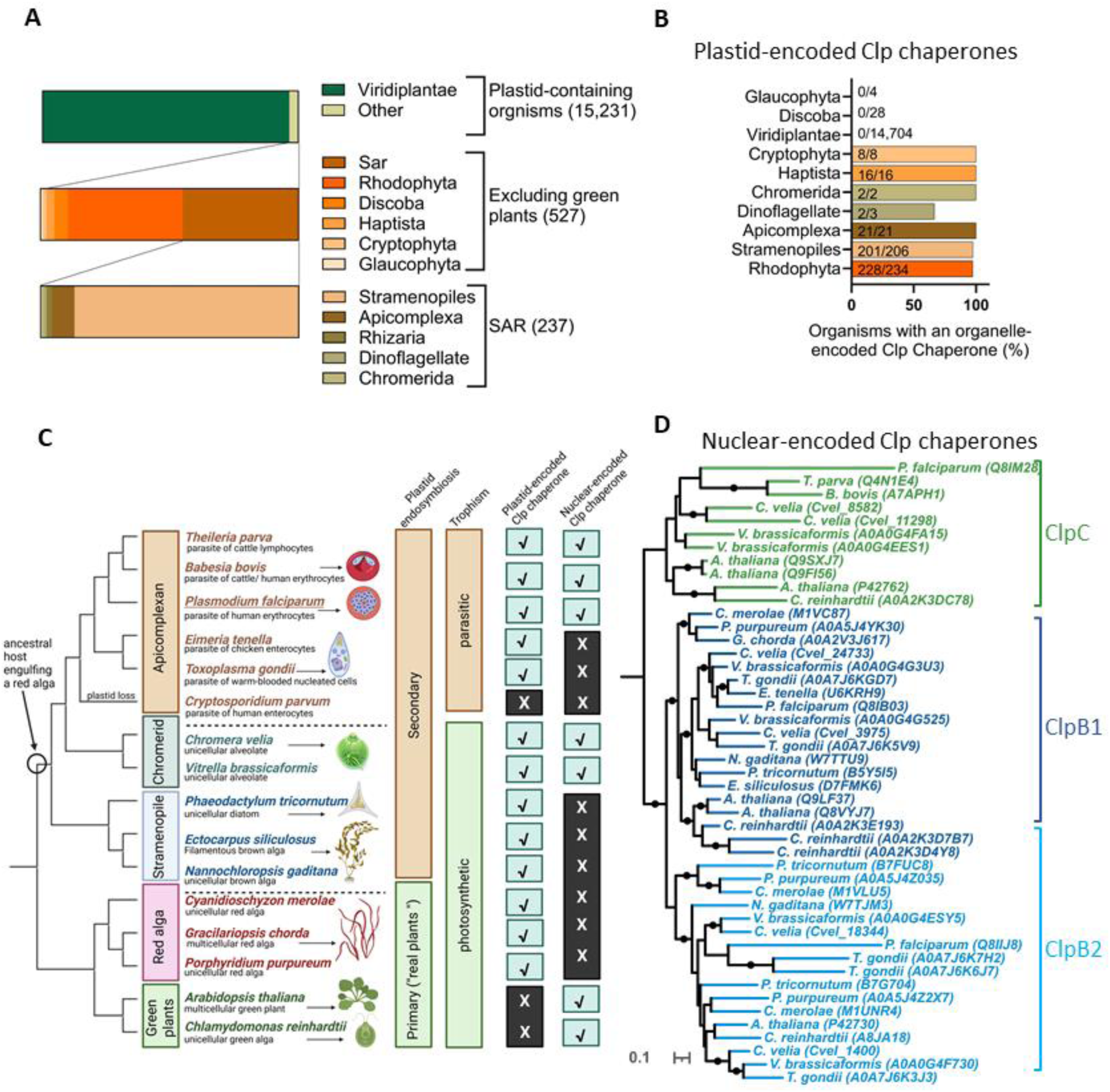
Phylogenetic analysis reveals conservation of plastid-encoded Clp chaperone. **(A)** Taxonomic distribution of plastid organelle genomes used in this study. A total of 15,231 complete plastid genomes were retrieved from the NCBI Organelle Genome Resources database. Genomes were classified into major taxonomic groups, including Viridiplantae (N = 14,704) and non-green plastid-containing lineages (N = 527). The latter includes Rhodophyta (red algae) and the SAR supergroup (N = 237), further subdivided into Stramenopiles, Apicomplexa, Rhizaria, Dinoflagellata, and Chromerida. Colours indicate major taxonomic groups. **(B)** Distribution of organelle-encoded *clpM* homologs across plastid-containing lineages. Predicted protein sequences from each plastid organelle genome were searched against the *Plasmodium falciparum clpM* sequence (PF3D7_API03600.1, 767 aa) using BLASTp (e-value ≤ 1e-5). The bar chart shows the percentage of organisms within each taxonomic group harbouring at least one organelle-encoded *clpM* homolog (numbers in the bars indicate the fraction of organisms with a hit). Organelle-encoded *clpM* homologs were detected exclusively in Rhodophyta and in lineages whose plastids derive from a secondary endosymbiosis of a red alga (Stramenopiles, Apicomplexa, Chromerida, Haptista, Cryptophyta, and Dinoflagellata), but were absent from lineages with plastids of non-red-algal origin (Viridiplantae, Glaucophyta, Discoba). **(C).** Species tree of 16 representative plastid-containing eukaryotes selected from the lineages identified in (A). The tree topology is based on the OpenTree synthetic tree^75^. The open circle marks the ancestral event of engulfing a red alga. Checkmarks and crosses indicate the presence or absence of a plastid-encoded Clp chaperone (based on B) and a nuclear-encoded Clp chaperone (based on D). *Cryptosporidium parvum,* which underwent complete plastid loss, lacks both. **(D)** Midpoint-rooted maximum likelihood phylogenetic tree of 47 nuclear-encoded Clp chaperone proteins from 15 representative plastid-bearing eukaryotes, generated using the LG+R5 model under IQ TREE (300 nonparametric bootstraps) and Bayesian inference (MrBayes(. Filled black circles indicate nodes with bootstrap support ≥ 90 and posterior probability ≥ 0.95. The tree reveals three well-supported monophyletic groups corresponding to ClpC (green), ClpB1 (light blue), ClpB2 (dark blue).

However, previous works in *Plasmodium* showed expression, function and essentiality of a nuclear-encoded Clp chaperone paralog^22,25^. Therefore, to look deeper into the pattern of Clp organelle-to-nucleus gene-transfer, we searched for Clp chaperone homologs encoded in nuclear genomes in plastid containing eukaryotes. Because complete nuclear genome sequences are unavailable for many of these organisms, we chose 16 representative organisms (Fig. 1C). Six of those are obligatory apicomplexan parasites, including *P. falciparum* and *Toxoplasma gondii*, all of which contain an apicoplast, besides *Cryptosporidium parvum* which had lost the entire organelle^34^. Apicomplexans belong to the higher clade of alveolates, of which two non-apicomplexan relatives were chosen; the unicellular marine photosynthetic Chromerids *Chromera* and *Vitrella*. Further removed but still within the SAR superphylum, three stramenopiles were chosen; The unicellular diatom *Phaeodactylum*, the unicellular brown alga *Nannochloropsis*, and the multicellular filamentous brown alga *Ectocarpus*, all are marine and photosynthetic. Finally, five ‘real plants’ were chosen; three are rhodophyte (i.e. red algae, both multicellular and unicellular), whose ancestral red alga has been hypothesized to be the endosymbiont that gave rise to the plastid in all the above-mentioned organisms^35^. The last two are green plants, and include the multicellular plant *Arabidopsis* and the unicellular green alga *Chlamydomonas* (Fig. 1C).

With the exception of the plastid-less *Cryptosporidium*, all of these organisms encode nuclear Clp chaperones, consistent with the well-documented roles of Clp complexes in plastid core functions (Fig. 1D). However, Clp chaperones can be further divided into sub-groups, and not all of them function similarly, nor are they direct paralogs of the plastid-encoded ClpM. Multiple sequence alignment enabled us to construct a phylogenic tree of the nuclear-encoded Clp chaperones, dividing them into three major groups; ClpB1, ClpB2 and ClpC (Fig. 1D). Out of these three subtypes, it is only ClpC that is predicted to bind the Clp protease and to function as an unfoldase mediating substrate degradation^36^. These functions and sequence similarities place together the nuclear-encoded ClpC and the plastid-encoded ClpM as closer homologs, compared to other Clp chaperones such as ClpB (Fig. S1). These nuclear-encoded ClpC orthologs were found in only three group of organisms; (1) *Plasmodium* and its two closest relatives *Babesia* and *Theileria*, (2) the photosynthetic chromerids *Chromera* and *Vitrella*, and (3) green plants (Fig. 1D). These are interpreted as three independent cases of ClpC/M nuclear transfer that resulted in two different scenarios; in green plants, the most distant clade in this group of organisms, an ancient nuclear transfer of ClpC completely replaced the chloroplast gene. However, in two separated cases of *Plasmodium* and chromerids, two copies, a nuclear and a plastid gene reside together in the same organism. Collectively, our analysis shows that the physical presence of the Clp chaperone within the red-alga derived plastids is driven by a fundamental constraint specific to this gene, and is potentially linked not only to protein homeostasis, but also to the retention of the plastid genome in these organisms. The two independent cases of nuclear-encoded Clp in apicomplexan and chromerids suggest that Clp nuclear expression is possible under certain conditions. These involve gene duplication and substantial changes in protein sequence, but it cannot replace the plastid-encoded homolog.

### ClpM expression profile corresponds to apicoplast development

The phylogenetic pattern of Clp chaperone conservation suggests a critical function rather than sequence retention of a pseudogene. To gain evidence for expression of the apicoplast-encoded Clp chaperone specifically in *P. falciparum* (previously termed ClpM^21^ and used here thereafter), we performed stage-specific transcript analysis across the intraerythrocytic-developmental cycle. Due to the high AT content of the coding gene (88.45%), measurements were highly variable and therefore we optimized various sets of primers, and used both quantitative real-time (qRT-PCR, Fig. 2A) and digital droplet (dd-PCR, Fig. 2B) as complementary measurement methods. The specific cell-cycle stages of these parasites were verified using blood smears (Fig. 2C). In both methodologies, expression levels of *clpM* remained low and stable throughout the ring, early and late trophozoite stages, and were markedly elevated at the schizont stage (Fig. 2A,B). To assess whether this expression pattern is associated with apicoplast biogenesis, we took parasites expressing apicoplast-targeted GFP and imaged them at the same corresponding stages in which mRNA was collected (Fig. 2D). These confirmed that endogenous ClpM levels peak together with apicoplast development, indicating stage-specific regulation during the intraerythrocytic developmental cycle.

**Figure 2.**
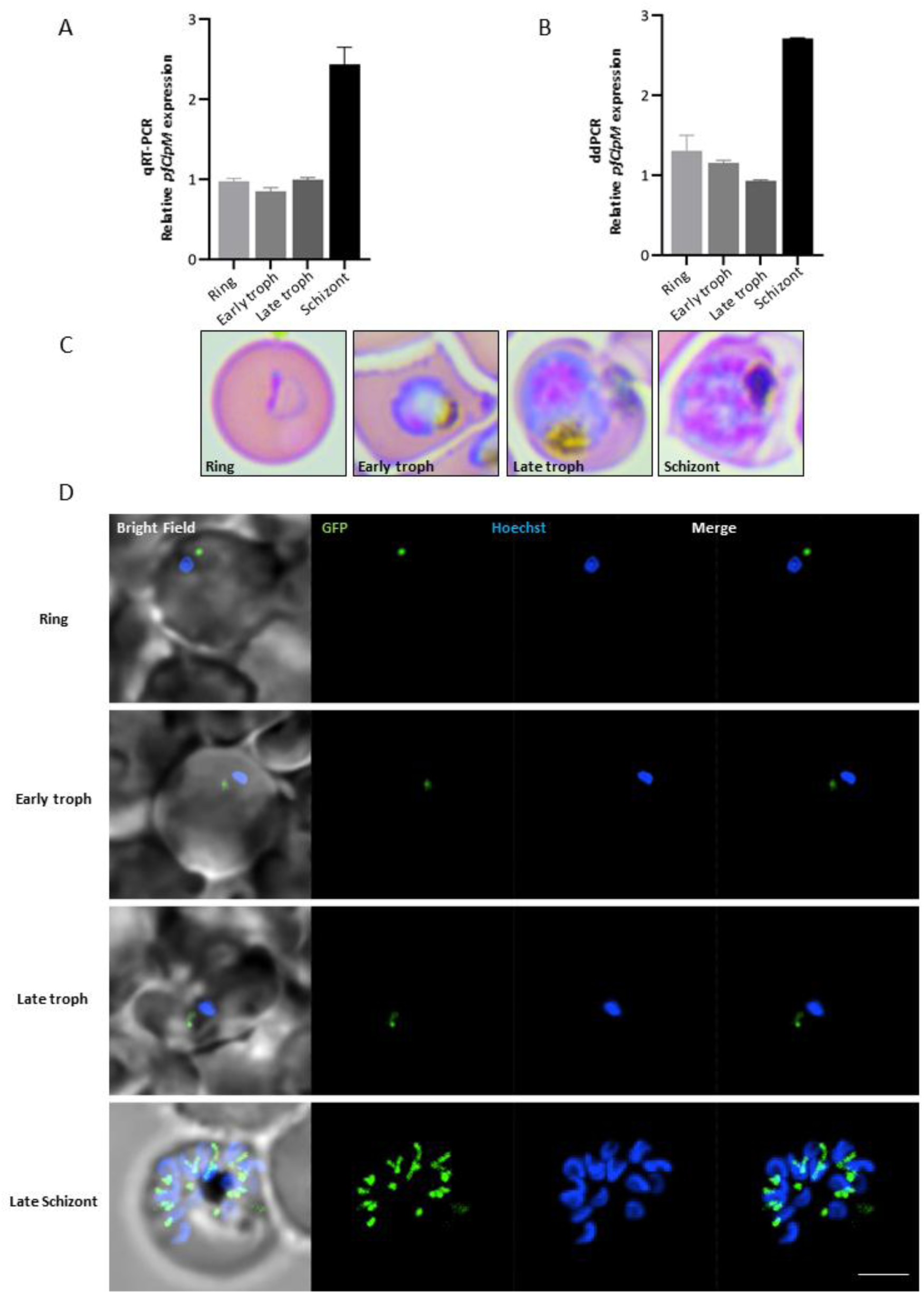
Stage-specific expression of endogenous *clpM* across the intraerythrocytic developmental cycle correlates with apicoplast development. **(A)** qPCR analysis of endogenous *clpM* transcript levels across parasite stages. Parasites were harvested from synchronized cultures at the indicated stages by saponin lysis, followed by RNA extraction and cDNA synthesis. Expression of *clpM* was normalized to aldolase. Data are shown from one representative experiment out of three independent biological replicates. Data are presented as mean ± SD of three technical replicates for each stage. **(B)** Same samples as in (A), analyzed by ddPCR. Error bars represent the mean of two independent primer sets targeting endogenous *clpM*. **(C)** Representative Giemsa-stained blood smears of a synchronized wildtype *P. falciparum* culture across one intraerythrocytic cycle, showing ring, early trophozoite, late trophozoite, and schizont stages. Images were acquired using an upright Eclipse E200 microscope. **(D)** Fluorescence microscopy of the indicated parasite stages. The apicoplast is marked by GFP (green), and nuclei are stained with Hoechst (blue). Images are shown as maximum intensity projections. Imaging was performed using a Nikon spinning disk confocal microscope equipped with a 100×/1.4 NA objective. Scale bar, 2 µm.

### Nuclear expression of ClpM using ectopic promoter results in a transcriptional block

The combination of a highly conserved genomic location with evident expression suggests that an evolutionary pressure specifically prevents the gene-transfer of *clpM* to the nucleus and retains it plastid-encoded. To test this hypothesis experimentally, we generated transgenic parasites expressing a second, nuclear-encoded copy of *clpM*. The nuclear-encoded *clpM* transgene was fused to an N-terminal apicoplast-targeting transit peptide (TP), and a C-terminal Ty tag (Fig. 3A). It was also fully recodonized, retaining the original amino acid sequence while increasing its GC content to facilitate cloning and nuclear expression and to distinct its transcript from the endogenous, plastid-encoded *clpM*. This design evaluates whether expression of *clpM* can occur outside its native genomic context or its localization and expression are evolutionarily constrained to the apicoplast.

**Figure 3:**
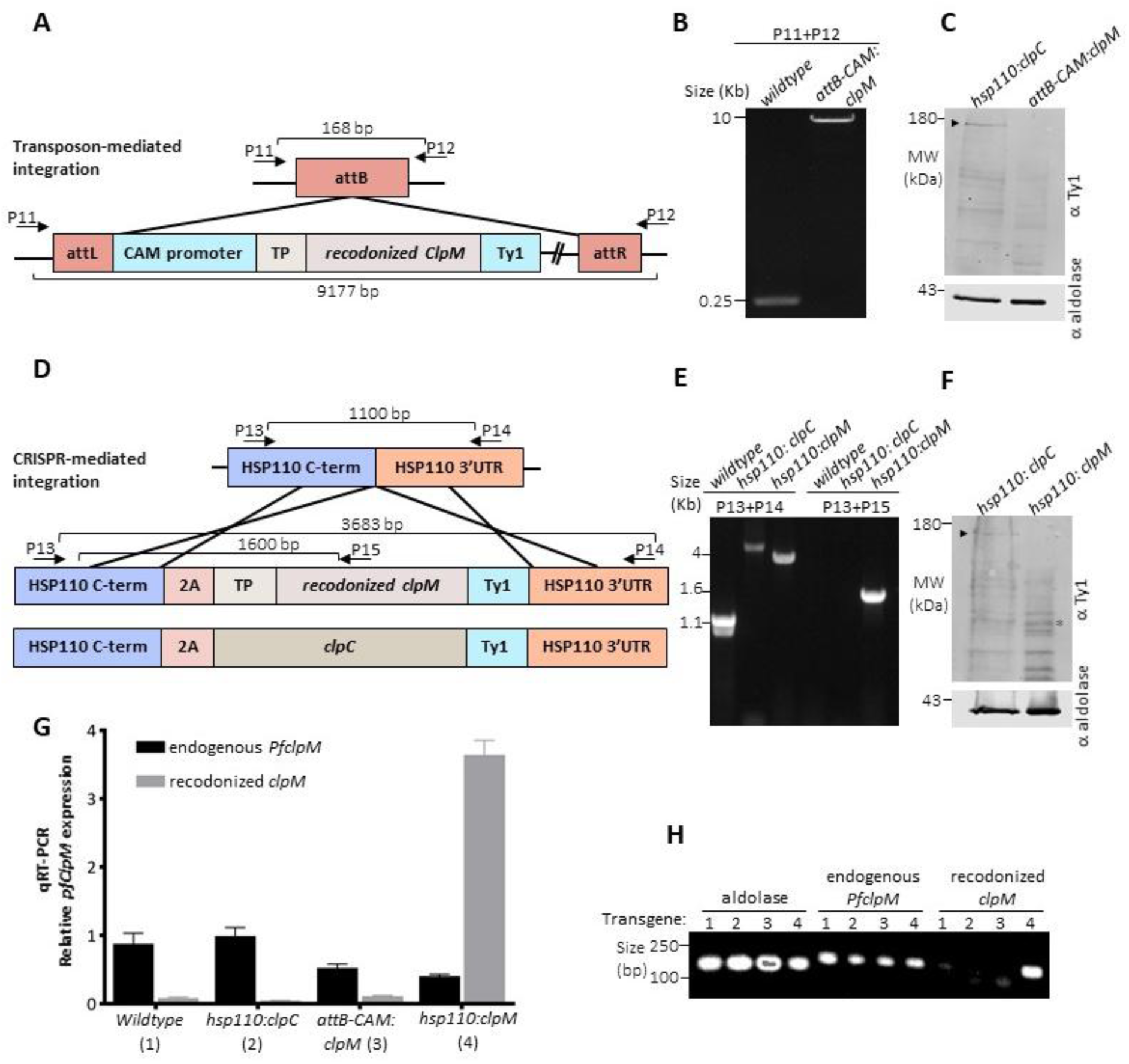
Expression analysis of nuclear-encoded clpM transgenes. **(A)** Schematic representation of the *attB-CAM:clpM* construct generated by BxB1-mediated integration into the parasite genome. **(B)** Genotyping PCR performed on gDNA confirming integration of the *attB-CAM:clpM* construct using P11+P12 primers. Amplification yields a 186 bp product in *wildtype* and a 9177 bp product in *attB-CAM:clpM* parasites. **(C)** Western blot analysis of lysates from *attB-CAM:clpM* parasites. Expression of the C-terminally Ty1-tagged ClpM protein (∼102 kDa) was probed using anti-Ty1 antibody; no signal was detected. ClpC (∼156 kDa, black arrowhead) served as a positive control. Aldolase (∼41 kDa) was used as a loading control. Equal amounts of protein were loaded per lane. **(D)** Schematic representation of the *hsp110:clpM* construct generated using CRISPR/Cas9-mediated integration at the *hsp110* locus. **(E)** Genotyping PCR performed on gDNA confirming integration of the *hsp110:clpM* construct. Amplification using P13+P14 primers yields products of 1100 bp (*wildtype*), 4087 bp (*hsp110:clpC*), and 3683 bp (*hsp110:clpM*). Amplification using P13+P15 primers yields a 1600 bp product exclusively in *hsp110:clpM* parasites. **(F)** Western blot analysis of lysates from *hsp110:clpM* parasites. Expression of the C-terminally Ty1-tagged clpM protein (∼102 kDa) was assessed using anti-Ty1 antibody; no signal was detected. ClpC (∼156 kDa) served as a positive control. Aldolase (∼41 kDa) was used as a loading control. Equal amounts of protein were loaded per lane. **(G)** qPCR analysis of transcript expression performed on cDNA derived from parasite RNA extracted from: (1) parental *NF54*, (2) *hsp110:ClpC,* (3) *attP-CAM:clpM,* and (4) *hsp110:ClpM*. *Endogenous pfclpM* served as a positive control, while expression of the *recodonized clpM* was assessed in transgenic lines. Arg-tRNA-ligase served as a reference gene for normalization. Data represent three independent biological replicates; error bars indicate SD of technical triplicates from a representative experiment. Equal amounts of cDNA were used across samples. **(H)** Agarose gel analysis of qPCR products. Aldolase was used as a reference gene and shows consistent amplification across all samples. *Endogenous pfclpM* is detected in all cell lines, whereas *recodonized clpM* is detected exclusively in *hsp110:clpM* parasites.

Our initial approach was to use the *attP-attB* transposon-mediated expression system, which we and others have successfully used in the past to generate multiple *Plasmodium* transgenes under the ubiquitous calmodulin promoter (*CAM*)^37,38^. Indeed, co-transfection of the *attP-CAM:clpM* plasmid with the Bxb1 integrase enabled full plasmid integration into the genomic *attB* locus, as confirmed by diagnostic PCR and Sanger sequencing (Fig. 3B). However, the *attB-CAM:clpM* transgene failed to produce any transcript (Fig. 3G) or protein (Fig. 3C). Because the *attB-CAM* system is typically streamlined^37,38^, a simple technical issue seemed unlikely. Instead, we suspected that expression of the *attB-CAM:clpM* transgene is associated with a toxic effect, which is overcome by transcriptional shutdown.

### Nuclear expression of ClpM using an endogenous promoter leads to translational block

To overcome this challenge, we designed a different strategy for ectopic expression that forces the parasite to transcribe the nuclear transgene. In this new strategy, we used CRISPR/Cas9 to integrate the recodonized *clpM* into an endogenous locus. The full *clpM* construct was integrated 3’ to the essential *hsp110* gene, separated by a 2A skip peptide. In this way, the endogenous *hsp110* promoter drives the co-expression of the native *hsp110* and transgenic *clpM* in a single transcript, avoiding transcriptional repression due to the essentiality of the *hsp110* gene (transgene termed *hsp110:ClpM,* Fig. 3D). As a control, we also expressed a second copy of the nuclear-encoded *clpC* under the *hsp110* expression system^22^ (transgene termed *hsp110:ClpC,* Fig. 3D). Accurate genomic integration was verified by PCR and Sanger sequencing (Fig. 3E). qPCR showed that this design produced high transcription levels of *hsp110:clpM* transgene whereas endogenous *ClpM* levels were unaltered (Fig. 3G). These results were further validated by visualization of amplification products on an agarose gel. Bands corresponding to aldolase and endogenous *clpM* were detected in all four cell lines; (1) parental *wildtype*, (2) *hsp110:ClpC,* (3) *attB-CAM:clpM,* and (4) *hsp110:ClpM*. In contrast, primers specific for transgenic recodonized *clpM* amplified a product only in *hsp110:ClpM*, further confirming that transcription of the nuclear-encoded transgene occurs exclusively in this engineered line (Fig. 3H). However, successful transcription did not lead to any significant protein expression of transgenic *hsp110:clpM* at the predicted molecular weight (Fig. S2). Western blot analysis showed successful protein expression of transgenic *hsp110:ClpC*, however *hsp110:clpM* remained undetectable at the protein level (Fig. 3F).

The sequence of *hsp110:clpM transcript* was confirmed by both Nanopore and Sanger sequencing and therefore we explored alternative reasons for the lack of protein expression. Our initial hypothesis was that *hsp110:clpM* protein is expressed, transported to the apicoplast owing to its attached TP, and is being actively degraded in the organelle. We therefore treated all the transgenes with chloramphenicol that blocks apicoplast translation and leads to organelle loss^39^, enabling us to detect accumulation of apicoplast proteins that would otherwise have been degraded in the organelle. Chloramphenicol treatment was supplemented with isopentenyl pyrophosphate (IPP), to keep the apicoplast-less parasites viable^8^ for continued nuclear-encoded proteins expression. However, although *hsp110:clpC* was expressed, this treatment did not lead to any accumulation of *hsp110:clpM* protein, indicating that it does not reach the apicoplast (Fig. 4A). Next, we hypothesized that, like other apicoplast-targeted proteins, translation of *hsp110:clpM* occurs at the rough ER, however it cannot get folded properly outside of the organelle and is being actively degraded by ER-associated degradation (ERAD) and the proteasome. To test this, we treated the cells for 8 hours with proteasome inhibitor MG132^40^, and tested whether accumulation of transgenic ClpM can be observed (Fig. 4B). However, we could not detect any signal, suggesting that ERAD or proteasomal degradation are not involved in transgenic ClpM processing. Additionally, we cultured the parasites for 24h in lower temperatures (34°C and 32°C) to increase total protein folding and stability^41^ but detected no signal, suggesting that folding is not the limiting step. Another possibility is that transgenic ClpM is expressed, but due to high hydrophobicity is embedded in ER membranes which are being discarded during protein extraction. To preclude this possibility, we omitted both boiling and sonication, retaining most membranes and yielding a soft, adhesive lysate suitable for SDS-PAGE analysis. While *hsp110:clpC* was successfully detected under these conditions, *hsp110:clpM* remained undetectable, indicating that membrane association does not account for the failure to detect transgenic ClpM (Fig. 4D). Altogether, these data suggest that the absence of detectable ClpM protein is not due to degradation, instability, or extraction artefacts, but instead reflects a fundamental block in translating ClpM when encoded in the nucleus. Moreover, these data imply that both transcriptional and post-transcriptional factors contribute to the apparent confinement of *clpM* expression to the apicoplast.

**Figure 4:**
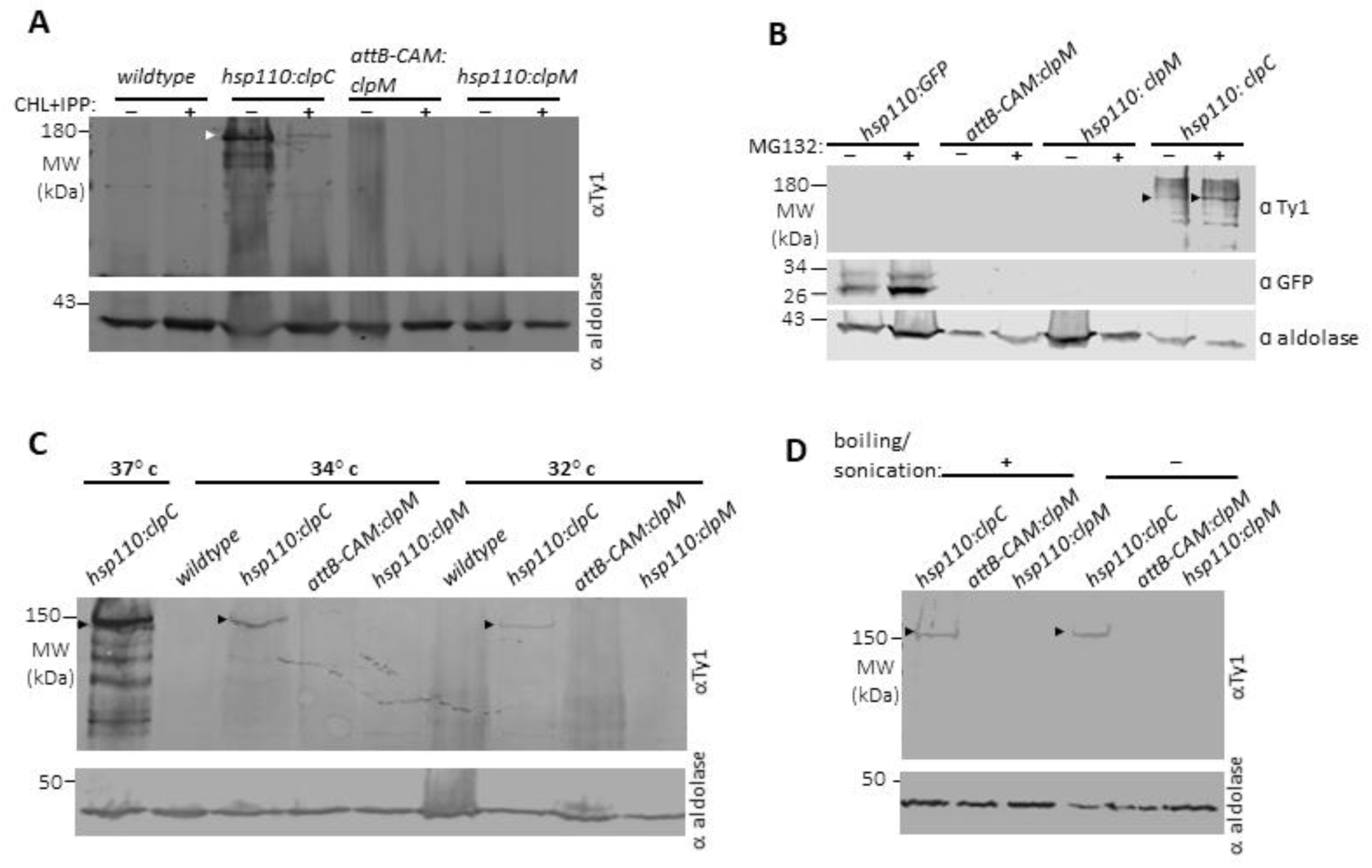
Assessment of transgenic ClpM protein expression and stability. **(A)** Western blot analysis of lysates from *wildtype* (negative control), *hsp110:clpC* (positive control), *attB-CAM:clpM*, and *hsp110:clpM* parasite lines grown in the presence of 40 µM chloramphenicol (CHL) and 200 µM isopentenyl pyrophosphate (IPP). Expression of C-terminally Ty1-tagged clpM (∼102 kDa) was assessed using anti-Ty1 antibody. No signal corresponding to ClpM was detected in *attB-CAM:clpM* or *hsp110:clpM* lines. ClpC (∼156 kDa, white arrowhead) was detected in the positive control. Aldolase (∼41 kDa) served as a loading control. **(B)** Western blot analysis of lysates from *wildtype*, *hsp110:clpC*, *attB-CAM:clpM*, and *hsp110:clpM* parasite lines treated with 200 nM MG132 for ∼6 h. Expression of Ty1-tagged proteins was assessed using anti-Ty1 antibody. No ClpM signal was detected in *attB-CAM:clpM* or *hsp110:clpM* lines. ClpC and GFP signals were detected in the positive controls (black arrowhead). Aldolase served as a loading control. Equal volumes of lysate were loaded per lane. **(C)** Western blot analysis of lysates from *wildtype*, *hsp110:clpC*, *attB-CAM:clpM*, and *hsp110:clpM* parasite lines cultured at 37°C, 34°C, or 32°C for 48 h. Expression of Ty1-tagged proteins was assessed using anti-Ty1 antibody. Expression of ClpC (black arrowhead) is visible, however no ClpM signal was detected under any temperature condition. Aldolase was used as a loading control. Equal volumes of lysate were loaded per lane. **(D)** Western blot analysis of lysates prepared using a membrane-preserving protocol (omitting sonication and boiling) from *hsp110:clpC*, *attB-CAM:clpM*, and *hsp110:clpM* parasite lines. Under these conditions, ClpC-Ty1 was detected (black arrowhead), whereas ClpM-Ty1 remained undetectable. Protein expression was probed using anti-Ty1 antibody, with aldolase as a loading control. Equal volumes of lysate were loaded per lane.

### Endogenous apicoplast nucleotide sequences lead to transgene toxicity

Our two reference transgenes were (1)*hsp110:clpC* which serves as a positive control and (2)*hsp110:clpM,* which is being transcribed but not translated. While the nucleic acid sequence of *hsp110:clpC* was identical to the endogenous nuclear gene, the sequence for *hsp110:clpM* was recodonized for the reasons outlined above. To explore the potential role of codon usage in organelle gene confinement, we amplified the original apicoplast-encode gene sequence to generate the construct *hsp110:clpM^endo^* (Fig. 5A). Similarly, we cloned another apicoplast-encoded gene with its original sequence, *SufB*, supposedly a key enzyme in the SUF pathway which builds essential iron-sulfur clusters in the apicoplast^12^. Transfections of both constructs, *hsp110:clpM^endo^* and *hsp110:SufB^endo^*, were carried out multiple times (42 and 26, respectively) and repeatedly failed to result in live parasites (Fig. 5B). This suggests that the original, non-recodonized sequences are non-viable when nuclear encoded, not only for ClpM but also for other apicoplast-encoded genes. Similarly, a ClpM construct with an additional N-terminal tag (*hsp110:clpM^x2Ty)^*, also failed to integrate over 30 attempts, supporting the potential cytotoxicity of the construct. Lastly, *hsp110:GFP*, a construct carrying a GFP reporter with a similar codon usage to the recodonized *hsp110:clpM*, was readily expressed (Fig. 5C). Collectively, the data indicate that apicoplast-encoded genes precluded from nuclear expression by at least two levels. Their nucleic acid sequence with an exceptionally low GC content renders them toxic while transcribed from the nucleus; and their amino acid sequence has a similar effect, leading to either transcription or translation block.

**Figure 5:**
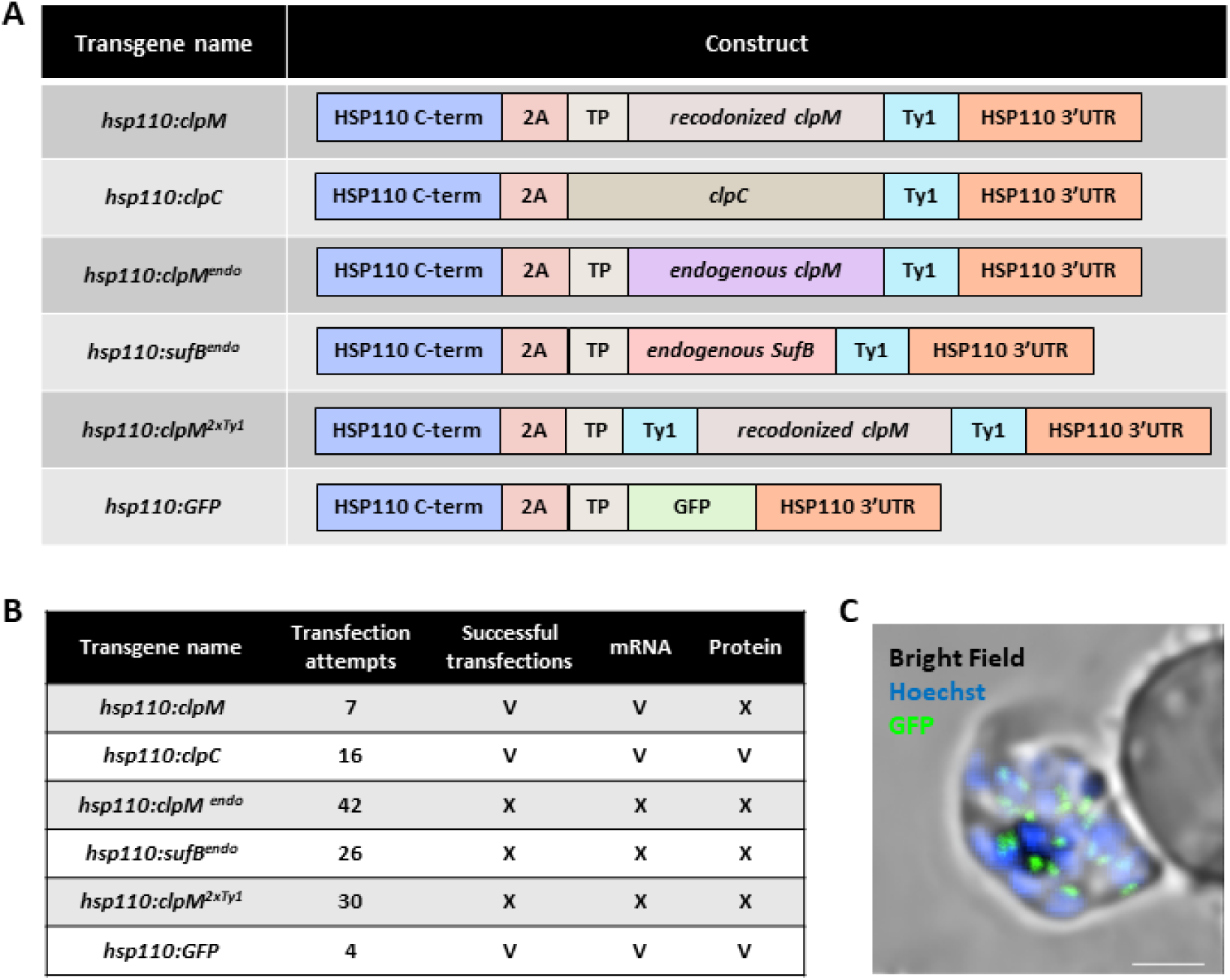
Generation and outcome of CRISPR/Cas9-mediated integration at the *hsp110* locus. **(A)** Schematic summary of constructs designed for integration at the *hsp110* locus in *Plasmodium falciparum* using CRISPR/Cas9. Constructs include *recodonized clpM* (*hsp110:clpM*), *clpC* (*hsp110:clpC*), *endogenous pfclpM* (*hsp110:clpM^endo^*), *sufB* (*hsp110:sufB^endo^*), *double Ty1-tagged recodonized clpM* with an additional N-terminal Ty1 tag (*hsp110:clpM^2×Ty^*), and *GFP* (*hsp110:GFP*). Integration of all constructs was done using the same strategy. **(B)** Summary of transfection outcomes. The number of transfection attempts and successful integrations is shown for each construct. *hsp110:clpC* and *hsp110:GFP* yielded successful integration with detectable mRNA and protein expression. *hsp110:clpM* showed successful integration and transcription but no protein expression. No successful integration was obtained for the remaining constructs. **(C)** Representative fluorescence live microscopy image of the *hsp110:GFP* transgenic parasite line. Image was captured as Z-stacks maximum intensity projection. GFP localizes to the apicoplast is green, and nucleus is labeled in Blue with Hoechst. The imaging was performed using a Nikon Spinning Disk confocal fluorescence microscope equipped with a 100x/1.4NA objective. Scale bar is 2.5 μM.

### Domain structure of ClpC/M chaperones is conserved across nuclear and plastid encoded orthologues

Within the SAR supergroup, the nuclear-encoded *clpC* gene emerged twice, in Apicomplexa and chromerids, but in both cases it did not lead to the loss of apicoplast-encoded *clpM* gene (Fig. 1D). Our transgenes demonstrate that expression of an additional nuclear copy was possible for ClpC but not ClpM, an experimental reflection of the evolutionary phenomenon (Fig. 3F). The facts that they are both expressed (Fig. 2), ClpC is essential^25^, and so is translation in the organelle^39^, collectively suggest that the two chaperones do not have completely redundant functions.

To look further into those possible distinct functions, we focused on the specific domains of ClpC/M orthologs. These mainly constitute two conserved AAA+ ATPases domains termed D1 and D2, and a ClpP binding motif, differentiating them from other Clp chaperones. In *P. falciparum*, both ClpC and ClpM contain D1 and D2 domains with 30.1% and 43.6% similarities respectively (Fig S1). However, in ClpM, the D1 Walker A motif is degenerate and the ClpP-binding site is absent, raising questions about whether canonical domain structure and function are conserved (Fig S1). We therefore analyzed all plastid- and nuclear-encoded ClpC homologs (as defined in Fig. 1) for sequence conservation and relationship between functional ATPases domains (Fig. 6A). This analysis revealed that D1 and D2 domains form separate clusters independent of genomic origin or species, suggesting that each domain fulfills conserved yet distinct functional roles across taxa. Moreover, in some plastid-encoded ClpM genes, including *Plasmodium*, the D1 domain is inactive and thus removed from analysis. These genes encode only a single functional D2 domain, suggesting that the D2 domain contains the most critical gene function (Fig. 6A).

**Figure 6:**
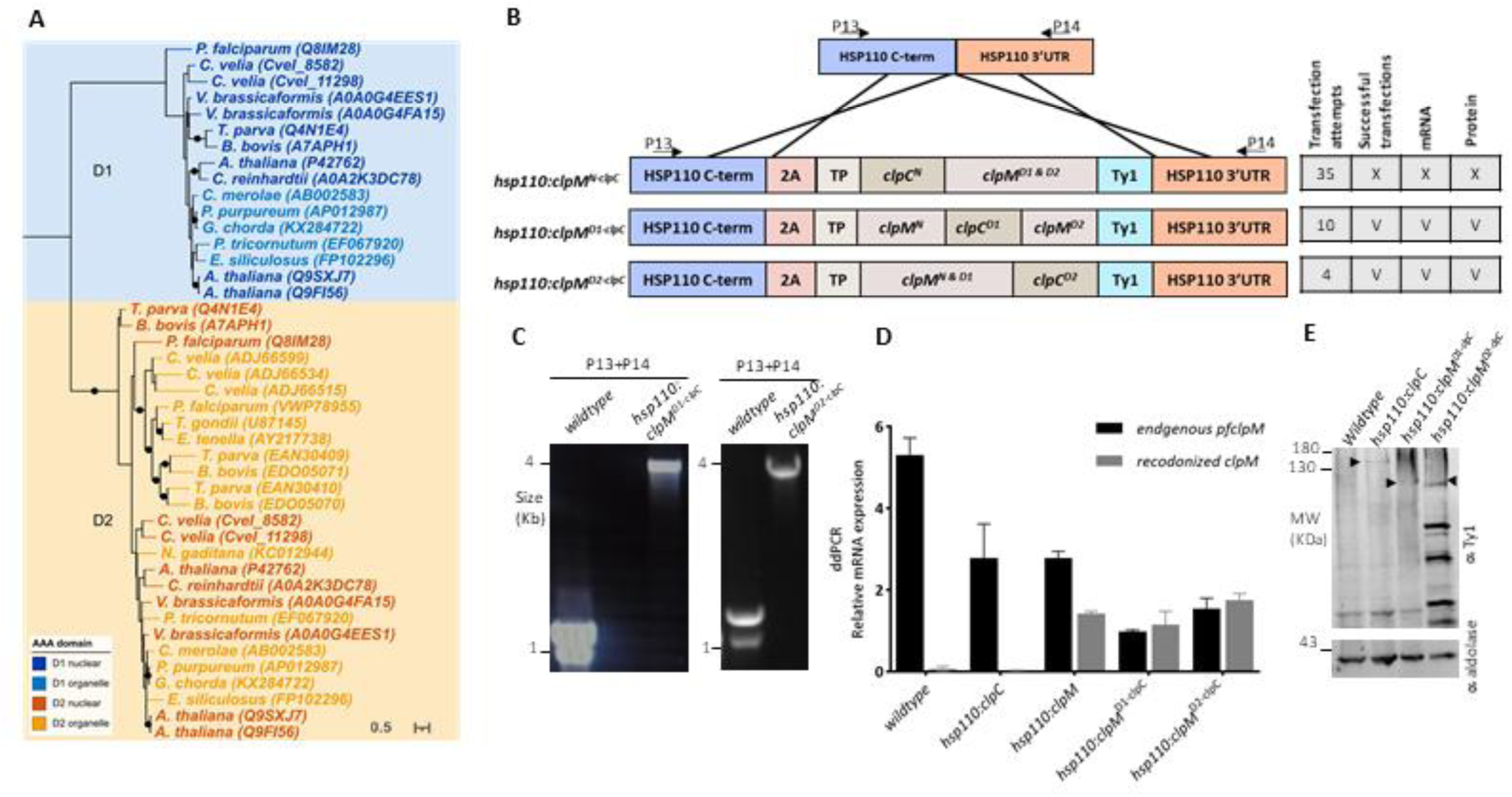
Phylogenetic and Expression analyses of ClpM domain swaps with ClpC. **(A)** Midpoint-rooted maximum likelihood phylogenetic tree of functional AAA ATPase domains from nuclear, and organelle encoded ClpC/M chaperones across 15 representative plastid-bearing eukaryotes, generated using the LG+I+G4 model under IQ-TREE (300 nonparametric bootstraps) and MrBayes (LG+I+G). Filled black circles indicate nodes with bootstrap support ≥ 85 and posterior probability ≥ 0.95. Each protein was split into its individual AAA ATPase domains (D1, D2), and both were included as separate sequences. The tree reveals a clear separation between D1 and D2 domains regardless of whether the protein is nuclear or organelle encoded, indicating that the two domains are functionally and evolutionarily distinct. Single-AAA-domain proteins cluster within the D2 clade, suggesting that when only one AAA domain is retained, it is invariably D2. Sequences are colored by domain and genomic origin: nuclear-encoded D1 (dark blue), organelle-encoded D1 (light blue), nuclear-encoded D2 (dark orange), organelle-encoded D2 (light orange). **(B)** Schematic representation of domain-swapped constructs generated at the *hsp110* locus and summary of transfection outcomes. Replacement of the N-terminal domain of clpM with that of clpC (*hsp110:clpM^N-clpC^*) did not yield successful integration. In contrast, replacement of either the D1 (*hsp110:clpM^D1-clpC^*) or D2 domain (*hsp110:clpM^D2-clpC^*) resulted in successful integration with detectable mRNA and protein expression. **(C)** Genotyping PCR performed on gDNA confirming correct integration of *hsp110:clpM^D1-clpC^* and *hsp110:clpM^D2-clpC^* parasite lines. **(D)** ddPCR analysis performed on cDNA derived from saponin-isolated parasites. Expression was assessed using two primer sets: one detecting *endogenous pfclpM* (detected in all lines, including *wildtype* and *hsp110:clpC*), and a second specific to the recodonized constructs (detected in *hsp110:clpM, hsp110:clpM^D1-clpC^* and *hsp110:clpM^D2-clpC^* lines). Equal amounts of cDNA were used across samples. Error bars indicate standard deviation (SD) of two primers sets for the same gene. One representative experiment is shown, out of three biological replicates. **(E)** Western blot analysis of lysates from saponin-isolated parasites following sonication. Equal amounts of protein were loaded per lane. Expression of Ty1-tagged proteins was detected using anti-Ty1 antibody. Bands corresponding to *hsp110:clpM^D1-clpC^* and *hsp110:clpM^D2-clpC^* are observed at the expected molecular weights (∼115 kDa and ∼114 kDa, respectively, black arrowheads). ClpC (∼156 kDa, black arrowhead) was detected in the positive control. Aldolase (∼41 kDa) served as a loading control.

### Domains swaps between ClpC and ClpM enable nuclear expression

We therefore decided to test methodically which domain of *clpM* gene confines it to the apicoplast genome. This strategy was led by the hypothesis that certain domains have been mutated enough in *clpC*, enabling its nuclear expression. Both genes contain the three domains expected of Clp chaperones: N-terminal domain, and two AAA+ domains with conserved Walker A and Walker B motifs^23^, and called here for convenience N, D1 and D2. Importantly, D1 of apicoplast-encoded *clpM* does not have the canonical Walker A and Walker B motifs and therefore is not expected to be a functional ATPase^21^. To test this hypothesis, we generated three additional constructs under the *hsp110* expression system, each a domain-swapped chimera between ClpM and ClpC (Fig. 6B). The *hsp110:clpM^N-clpC^*construct, in which the N-terminal domain of ClpM was replaced with the corresponding domain from ClpC, failed to produce any successful transfection despite 35 attempts (Fig. 6B). This indicates that the N-terminal domain of ClpM is not the source hindrance, as its swapping did not improve nuclear expression.

In contrast, both the *hsp110:clpM^D1-clpC^* and *hsp110:clpM^D2-clpC^* chimeras (having D1 or D2 domains of ClpM replaced with the corresponding domain from ClpC) resulted in successful integrations (Fig. 6C). In both transgenes, a viable transcript was detected both via ddPCR (Fig. 6D) and qPCR (Fig. S3). Importantly, in contrast to all other ClpM constructs, the *hsp110:clpM^D1-clpC^* and *hsp110:clpM^D2-clpC^*chimeras resulted in a detectable protein expression in the corresponding band size (115 & 114 kDa respectively, Fig. S2), in addition to smaller bands, potentially representing the more stable domains (Fig. 6E). This suggests that the cytotoxicity/ translation block of nuclear ClpM can be traced to one of its conserved AAA+ domains, as replacing either one with the ClpC equivalent enables partial expression.

### ClpM-ClpC chimeras are degraded in the ER and are associated with a fitness cost

We next investigated whether the ClpM-ClpC chimeras have any effect on apicoplast function and specifically translation, with the rationale being that a key organelle gene transported from the outside may render its organellar translation obsolete. To do that, we tested whether sensitivity to chloramphenicol, the apicoplast ribosome inhibitor, is altered in any of the transgenes. However, the half-maximal effective concentration (EC50) for Chloramphenicol was comparable in parental *wildtype* strain (6.06mM), *hsp110:clpM^D1-clpC^* (3.54mM) and *hsp110:clpM^D2-clpC^* (4.49mM) (Fig. 7A). We therefore investigated whether the ClpM-ClpC chimeras transgenes reach the apicoplast, and getting partially degraded in the organelle. To do that, we treated transgenes with chloramphenicol for apicoplast ablation, supplemented with IPP to maintain viability, and extracted protein to detect any accumulation or modified degradation pattern. The *hsp110:GFP* control demonstrated the expected pattern of a double band in untreated parasites, consisting of a larger band of the full-length protein, and a smaller one corresponding to the main apicoplast-localized fraction, post-removal of the transit peptide (Fig. 7B, left). Upon treatment, an accumulation of the upper band is observed, consistent with organelle loss (Fig. 7B, left). However, this was not observed with either *hsp110:clpM^D1-clpC^* or *hsp110:clpM^D2-clpC^*, both showing very weak and unstable expression regardless of treatment, suggesting they do not reach the apicoplast (Fig. 7B, right). To validate this observation, we visualized the transgenes using a combination of live and immunofluorescence microscopy. Using live imaging, we could readily detect the typical apicoplast morphology in the *hsp110:GFP* transgene (Fig. 7C). Likewise, using immunofluorescence assay (IFA) with an anti-Ty antibody, we could detect the typical elongated apicoplast form in the *hsp110:ClpC* transgene (Fig. 7D). Consistent with no signal on a Western Blot (Fig. 3F), the *hsp110:ClpM* transgene did not produce any IFA signal (Fig. 7E). However, both *hsp110:clpM^D1-clpC^* and *hsp110:clpM^D2-clpC^* chimera transgenes showed diffuse signals which were markedly distinct from an apicoplast localization, and their perinuclear positioning suggested an ER localization (Fig. 7F & 7G).

**Figure 7:**
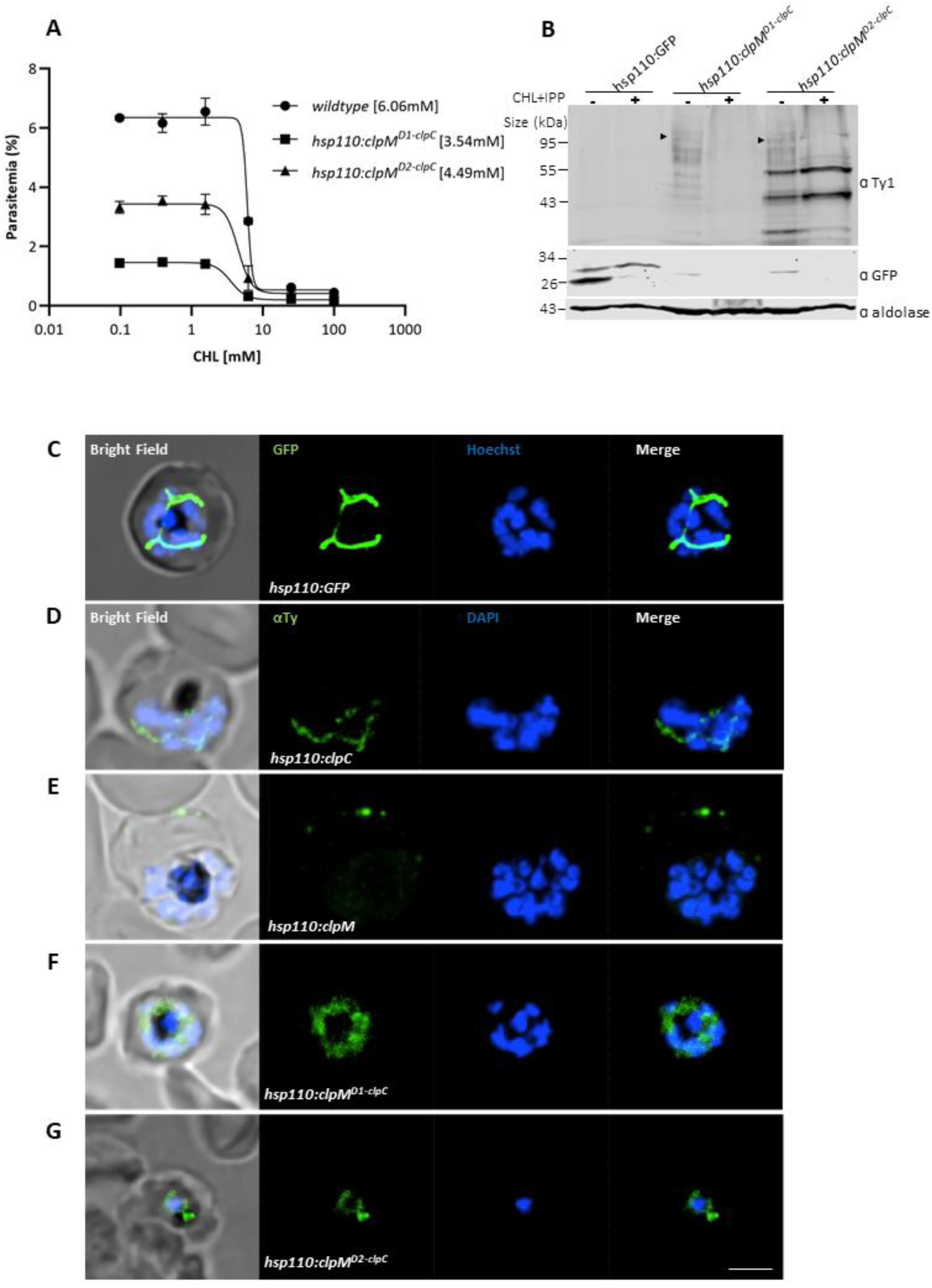
Localization analysis of ClpM-ClpC domain-swap parasites. **(A)** Dose–response analysis of parasite growth following CHL treatment. *Wildtype*, *hsp110:clpM^D1-clpC^*, and *hsp110:clpM^D2-clpC^* parasite lines were incubated with serial dilutions of CHL in 96-well plates. Parasitemia was measured after 120 h by flow cytometry. EC50 values were calculated as 6.06 nM (*wildtype*), 3.54 nM (*hsp110:clpM^D1-clpC^*), and 4.49 nM (*hsp110:clpM^D2-clpC^*). Data were fitted to a dose–response curve and are presented as mean ± SEM. **(B)** Western blot analysis of lysates from *hsp110:GFP* (positive control), *hsp110:clpM^D1-clpC^*, and *hsp110:clpM^D2-clpC^*parasite lines cultured in the presence of 40 µM CHL+200 µM IPP. Expression of C-terminally Ty1-tagged proteins was assessed using anti-Ty1 antibody. Bands corresponding to *hsp110:clpM^D1-clpC^* (∼115 kDa) and *hsp110:clpM^D2-clpC^*(∼114 kDa) were evaluated. Weak signal was detected for *hsp110:clpM^D1-clpC^*, whereas *hsp110:clpM^D2-clpC^* showed detectable expression with no apparent change upon CHL+IPP treatment. In *hsp110:GFP* parasites, was detected using anti-GFP antibody. In untreated conditions, two forms were observed: a higher molecular weight band corresponding to TP-GFP (∼34 kDa) and a lower band corresponding to processed GFP (∼27 kDa). Upon CHL treatment, only the higher molecular weight form was detected. Aldolase (∼41 kDa) served as a loading control. Equal amounts of protein were loaded per lane. **(C-G)** Fluorescence microscopy analysis of transgenic parasite lines. Images were captured as Z-stacks maximum intensity projection. Imaging was performed using a Nikon Spinning Disk confocal fluorescence microscope equipped with a 100x/1.4NA objective. Scale bar is 2 μM. (C) Fluorescence live microscopy image of the *hsp110:GFP* transgenic parasite line. GFP localizes to the apicoplast is green, and nucleus is labeled in Blue with Hoechst. (D-G) Immunofluorescence assay (IFA) with anti-Ty1 antibody (green) and DAPI (blue) for: **(D)** *hsp110:clpC,* **(E)** *hsp110:clpM,* **(F)** *hsp110:clpM^D1-clpC^*, and **(G)** *hsp110:clpM^D2-clpC^*transgenes.

To verify this subcellular localization as well as to test for potential reasons for the ClpM-ClpC chimeras increased instability, we grew parasites in low temperatures as well as treated cultures with proteasome inhibitor MG132. While low temperatures did not have any stabilizing effect (Fig. 8A), inhibition of proteasomal degradation markedly reduced degradation bands and mildly increased the full-length product of the transgenes at their predicted molecular weights (Fig. 8B). This, together with the microscopy data (Fig. 7C), suggest a scenario in which the *hsp110:clpM^D1-clpC^* and *hsp110:clpM^D2-clpC^* chimera transgenes are being co-translationally imported to the ER as expected due to their transit peptide. In the ER, they fail to properly fold and are being recognized by the ERAD machinery, and subsequently degraded by the proteasome.

**Figure 8.**
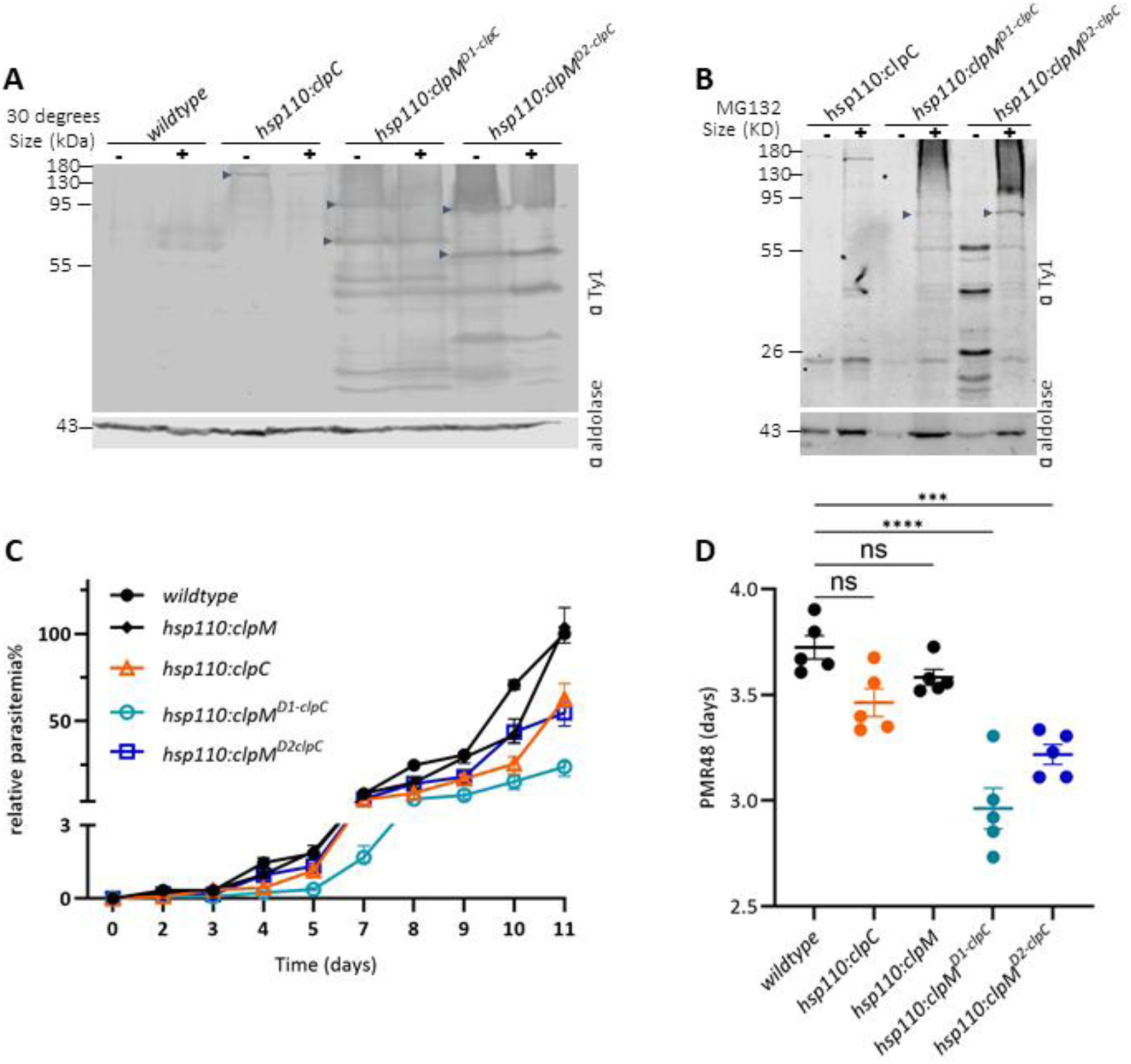
ClpM-ClpC chimeras are degraded in the ER and are associated with a fitness cost. (A) Western blot analysis of lysates from *wildtype*, *hsp110:clpC*, *hsp110:clpM^D1-clpC^*, and *hsp110:clpM^D2-clpC^* parasite lines cultured at either 37°C (-) or 30°C (+) for 48 h. Expression of transgenic proteins was assessed using anti-Ty1 antibody. Expression of ClpC (∼156 kDa; black arrowhead) was detected in the positive control, and multiple ClpM chimeras bands were detected under both temperature conditions. Aldolase was used as a loading control. Equal volumes of lysate were loaded per lane. **(B)** Western blot analysis of lysates from *hsp110:clpC*, *hsp110:clpM^D1-clpC^*, and *hsp110:clpM^D2-clpC^* parasite lines treated with 200 nM MG132 for ∼6 h. Expression of Ty1-tagged proteins was assessed using anti-Ty1 antibody. ClpM bands corresponding to the expected molecular weights were detected in both *hsp110:clpM^D1-clpC^*and *hsp110:clpM^D2-clpC^* lines. ClpC (∼156 kDa; arrowhead) was detected in the positive control. Aldolase served as a loading control. Equal volumes of lysate were loaded per lane. **(C)** Growth curve of *wildtype*, *hsp110:clpC*, *hsp110:clpM, hsp110:clpM^D1-clpC^*, and *hsp110:clpM^D2-clpC^* parasites. Parasitemia was monitored every 24h over 11 days via flow cytometry. Parasitemia of the *wildtype* strain at the end of each experiment was set as the highest relative parasitemia and was used to normalize parasites growth. Normalized data are represented as mean ± SEM for four technical replicates. **(D)** Parasitemia of *wildtype*, *hsp110:clpC*, *hsp110:clpM, hsp110:clpM^D1-clpC^*, and *hsp110:clpM^D2-clpC^* parasite lines was measured over time by flow cytometry. Data were log₁₀-transformed and plotted against time (days), and linear regression was applied to estimate parasite growth rates (m), assuming exponential growth. The parasite multiplication rate over 48 hours (PMR₄₈) was calculated using the equation PMR₄₈ = 10^(2m), where m represents the slope of the log₁₀-transformed parasitemia. Data are presented as mean ± SEM from n = 5 biological replicates. Statistical significance was determined using one-way ANOVA followed by multiple comparisons relative to the *wildtype* control.

Finally, we were interested to see whether any of the transgene is associated with a fitness cost that would manifest itself in the form of compromised growth. To do that, the various transgenes were seeded and allowed to grow for 11 days while parasitemia was monitored daily by flow cytometry. The *hsp110:ClpM* transgene grew at the same rate as the parental *wildtype* strain, consistent with the fact that there is no protein expression (Fig. 8C, black curves). The *hsp110:ClpC* and the *ClpM^D2-ClpC^* chimera transgenes grew at comparable rates, slightly slower than *wildtype* (Fig. 8C, blue and orange curves). The common domain of these two transgenes is the D2 domain of ClpC, which seems to be the most stable region in the *hsp110:clpM^D2-ClpC^* chimera (Fig. 8B, 50 kDa, asterisk), and thus may lead to the slower growth in both transgenes. Finally, *ClpM^D1-ClpC^* exhibited the slowest growth (Fig. 8C, teal curve) as could also be reflected in the longest replication time (Fig. 8D), potentially attributed to be carrying the functional D2 domain of ClpM.

## Discussion

Endosymbiont-to-organelle evolution is characterized by extensive gene loss and gene transfer from the organelle to the nucleus, yet this process repeatedly stalls with a small set of genes that remain organelle-encoded. While the logic of retaining photosynthesis-linked genes in chloroplast genomes has been debated for decades^18^, these photosynthesis-centric arguments do not explain why heterotrophic plastids, including the apicoplast of *apicomplexan* parasites, also retain a genome. In this work, the evolutionary “sticking point” was interrogated experimentally by attempting to force nuclear relocation of a candidate retention gene, *clpM*, and by dissecting the molecular barriers that prevent its nuclear expression. Collectively, the data support a model in which plastid gene retention can reflect not only selective advantages of organellar expression, but also mechanistic incompatibilities that block nuclear expression at multiple levels.

### ClpM is plastid-confined across lineages and shows developmental regulation

A first key observation is that clpM orthologues remain plastid-encoded across *Plasmodium*-related and ancestral lineages, whereas other Clp-family components are nuclear-encoded and targeted to the plastid/apicoplast (Fig. 1). This gene-specific confinement suggests evolutionary constraints acting particularly on ClpM rather than on the Clp system as a whole. Moreover, we found that the gene is highly conserved in many distant clades carrying rhodophyte-related plastids. This goes in line with multiple recent studies using specifically the sequence of the *clpM* gene as a genetic marker to determine phylogeny of *Plasmodium* infections in humans^42^ and primates^43^, *Haemosporidian* parasites in bats^44^, and newly-described apicomplexan endangering Japanese sea-cucumbers^45^. In parallel, stage-resolved transcript measurements indicate that endogenous *clpM* transcript abundance increases sharply at the schizont stage, consistent with a regulated role aligned with organelle growth and division. Although the physiological function of plastid-encoded ClpM in *Plasmodium* remains unclear, the combination of evolutionary conservation and stage-dependent transcription supports the idea that ClpM participates in a time-sensitive apicoplast process (e.g. organelle biogenesis) rather than representing a dispensable genomic relic.

These findings resonate with broader work establishing the importance of the apicoplast Clp network. El Bakkouri and colleagues catalogued multiple parasite Clp ATPases and protease components, distinguishing the plastid-encoded ClpM (historically annotated as ClpC in apicoplast genomes) from nuclear-encoded Clp chaperones^21,24^. Moreover, genetic studies have shown that nuclear-encoded apicoplast Clp components are essential for organelle integrity, underscoring the centrality of Clp-mediated proteostasis in apicoplast maintenance^22,25,46^. Together, these prior studies provide an important framework: Clp biology is essential in the apicoplast, making the persistent plastid encoding of *clpM* especially intriguing.

### Nuclear transfer is blocked by transcriptional silencing as well as translation barrier

The experimental strategy explicitly tested whether ClpM can be functionally expressed from the nucleus when supplied with a canonical apicoplast targeting peptide and (in the first iteration) codon recoding to facilitate nuclear expression. Two complementary outcomes were observed, corresponding to two levels of barrier. First, attB-mediated integration under the calmodulin promoter produced correct genomic integration but no detectable transcript or protein (Fig. 3), consistent with a parasite response of transcriptional shutdown, which is interpreted as an adaptive silencing of a toxic transgene. Second, to bypass promoter-level suppression, *clpM* was placed under enforced transcription by integrating it downstream of the essential *hsp110* locus via a 2A strategy. This design yielded robust *clpM* mRNA, verified by qPCR, ddPCR and sequencing, yet still no detectable ClpM protein, while a matched *hsp110:clpC* control was readily expressed (Fig. 3). The downstream experiments (apicoplast loss; proteasome inhibition; reduced-temperature; and altered lysis conditions) collectively argue against simple explanations such as apicoplast-localized degradation, ERAD clearance, generalized misfolding, or extraction artefacts (Fig. 4). The most parsimonious interpretation is that nuclear-encoded *clpM* is subject to a fundamental block at or before translation. Importantly, this is not a technical or generic limitation of apicoplast-targeted proteins or of AAA+ ATPases: the nuclear ClpC and GFP transgene controls could be expressed in the same framework.

Mechanistically, several (non-mutually exclusive) routes could explain a “translation barrier” for a plastid-encoded gene when expressed on ER-associated ribosomes: (i) sequence features in the mRNA that trigger ribosome stalling; (ii) co-translational quality control that rapidly clears nascent chains that form aberrant structures; or (iii) a requirement for organelle-localized translation to ensure immediate capture by specific chaperones/assembly factors. The HSP100/Clp family is built around conserved AAA+ ATPase modules that remodel higher-order protein assemblies, and subtle changes in these domains can alter interaction landscapes and conformational cycles^23^. This makes it plausible that a protein evolved for plastid translation and folding environments may be “non-permissive” when synthesized in the cytosol/ER-proximal context, even if the final destination is the apicoplast. Indeed, a recent study showed that apicoplast-targeted proteins involved in non-coding-RNA processing, indirectly affect translation of apicoplast-based transcripts, including *clpM*^47^.

### Two separable constraints: nucleotide-sequence toxicity and protein-intrinsic incompatibility

A particularly informative aspect of the study is the comparison between recodonized *clpM* and “endogenous-sequence” constructs. When the native apicoplast coding sequence was placed under the *hsp110* system (*hsp110:clpM^endo^*), repeated transfections failed to recover viable parasites; similarly, repeated attempts to express another plastid gene (*sufB*) from the nucleus also failed. These results support the conclusion that plastid-derived nucleotide sequences themselves can be toxic or otherwise incompatible with nuclear expression, likely reflecting extreme base composition and the presence of sequence motifs that disrupt nuclear transcription/processing (e.g., cryptic termination/polyadenylation signals, or RNA processing defects). In contrast, recoding enabled stable transcription of *clpM* in the nucleus, indicating that at least some nucleotide-level constraints can be bypassed. Yet, protein expression still did not occur. This combination supports a “two-layer” model:

1. Nucleotide constraints strongly disfavour nuclear expression of native plastid genes.
2. Protein-intrinsic determinants can impose an additional, post-transcriptional barrier even after mRNA is successfully produced.

### Domain swaps map the nuclear-expression barrier to the AAA+ D1/D2 modules

To localize the protein-level incompatibility, domain-swapped chimeras were engineered between ClpM and the nuclear ClpC. Replacement of the N-terminal region of ClpM (ClpM^N-ClpC^) did not “rescue” expression and in fact did not yield successful transfectants despite extensive attempts. In contrast, chimeras replacing either the D1 or D2 AAA+ domains of ClpM with the corresponding ClpC domains, generated transgenic parasites and produced detectable transcripts with partial translational recovery. These outcomes argue that the D1/D2 domains contribute substantially to the nuclear-expression incompatibility, whereas the N-terminus is not the dominant determinant.

This mapping is conceptually important for plastid genome evolution. It suggests that organelle-to-nucleus relocation of a gene like *clpM* may require adaptive changes in core functional domains, not only changes in targeting peptides or codon composition. In other words, nuclear relocation may be gated by the need to evolve a functional version of the protein that can be synthesized and handled by cytosolic translation/quality-control systems, representing an evolutionary hurdle that could preserve plastid encoding genes.

### Integrating these findings with hypotheses for plastid genome retention

Several ideas have been advanced to explain why plastid genomes persist in non-photosynthetic lineages. Two influential hypotheses were previously articulated^48^: (i) the “essential tRNAs” idea, and (ii) the “limited transfer window” hypothesis. As its name implies, the “essential tRNAs” hypothesis states that specific plastid tRNAs cannot be replaced/imported, which is plausible, but doesn’t explain why a translation machinery needs to be retained at the first place. Conversely, the “limited transfer window” hypothesis argues that gene transfer requires leakage of plastid DNA into the cytoplasm, and the frequency of such events is constrained in organisms with very few plastids per cell, where organelle lysis would be lethal. A key implication of the transfer-window model is that gene transfer is limited by opportunity, not necessarily by functional incompatibility.

The present work raises a third hypothesis; functional incompatibility is a major component of the barrier. At least for ClpM, even when nuclear transcription is enforced and a correct mRNA is produced, translation is blocked. This implies that a subset of apicoplast-encoded genes remains intrinsically non-permissive to nuclear expression, regardless of opportunities for DNA transfer or specialized tRNAs.

This phenomenon resonates more with the hydrophobicity hypothesis for mitochondrial genome retention. It argues that some genes are mitochondrial encoded because they are too hydrophobic for trafficking, and their cytosolic expression may result in mis-localization and cytotoxicity. This could be an attractive explanation for apicoplast genome retention as it prioritizes protein intrinsic properties. However, Kyte-Doolittle analysis reveals that ClpM is not particularly hydrophobic, and its hydrophobicity levels are comparable to the nuclear ClpC, arguing against such explanation (Fig. S4).

### Relationship to previous models: sufB/clpC as “gatekeeper genes” in Apicomplexans

Janouškovec and colleagues proposed a parsimonious model for plastid retention in Apicomplexans and their distant relatives, collectively referred to as Myzozoans^20^. According to that model, plastid genome loss becomes feasible only when a very small set of key plastid genes, particularly *sufB* and *clpC*, are transferred to the nucleus. The current data provide experimental texture to this evolutionary narrative. First, repeated failure to recover parasites expressing nuclear *clpM* and *sufB* with their original nucleotide sequences supports the idea that at least some “gatekeeper” plastid genes may be difficult to relocate because of sequence-level toxicity/incompatibility. Second, the *clpC/clpM* nomenclature problem is clarified experimentally: in *Plasmodium*, a nuclear *clpC* exists and is expressible, whereas the plastid-encoded *clpM* is the one that is stubbornly plastid-confined. The domain-swap results further suggest how the “relocation becomes feasible” theory might occur in principle: successful nuclear relocation may require evolutionary remodelling of AAA+ domains to render the protein permissive for cytosolic translation, consistent with the existence of nuclear *clpC* genes in some lineages, including *Plasmodium*.

### Limitations and Conclusions

A substantial limitation of this work is that the translation block is inferred from repeated failure to detect protein and from perturbations that did not unmask it; Moreover, the precise molecular mechanism (e.g. ribosome stalling versus rapid co-translational clearance) remains unresolved. Experimentally, this is addressed by a plethora of (1) alternative strategies (different promoters and integration systems, various codon usage), (2) positive controls (ectopic expressions of GFP and ClpC) and (3) partial expression rescue of D1 and D2 chimeras. Conceptually, these ‘negative results’ are an accurate experimental representation of the evolutionary failure to complete gene-transfer. Future studies with other plastid genes and different organisms will assist with generalizing these conclusions beyond *Plasmodium* and its relict plastid.

In sum, the results support an evolutionary model in which apicoplast genome retention is reinforced by mechanistic barriers to nuclear expression, rather than being explained solely by advantages of organellar control. ClpM emerges as a prime example of such a barrier: its nuclear expression is countered by transcriptional silencing/toxicity, and protein-intrinsic translation incompatibility mapping largely to the D1/D2 AAA+ domains. In the broader context, these findings give experimental grounding to models that emphasize a small number of plastid-encoded “gatekeeper” genes, while also expanding them by showing that “transfer feasibility” may require deep molecular adaptation of both nucleotide and protein features.

## Materials and Methods

### Plasmid construction

Genomic DNA from *Plasmodium falciparum* was isolated using the NucleoSpin Blood kit (Macherey-Nagel). PCR products were cloned into the appropriate plasmid backbones using In-Fusion Snap Assembly Master Mix (Takara). Restriction enzymes were purchased from New England Biolabs. All constructs were verified by Sanger sequencing (Hylabs). Plasmids were amplified by transformation into competent Stellar *E. coli* cells (New England Biolabs GmbH), cultured overnight at 37°C, and purified using the NucleoBond Xtra Midi Kit (Macherey-Nagel).

Oligonucleotides used in this study are summarized in **Table 1**. To clone plasmid pLN-ClpM^Ty^ (Full length, recodonized), the *PfclpM* gene (PF3D7_API03600) was recodonized using a codon usage adequate for *E. Coli* (Twist), fused to a N-terminal ACP transit peptide and C-terminal tripl-Ty1 tag. The synthesized fused gene was then amplified and cloned into pLN plasmid^37^ using primers **P5 + P6.** The construct was Sanger-sequenced using primers **P7 + P8 +P9 + P10** and inserted into pLN plasmid to produce *attB-CAM:clpM* construct, in which expression is driven by the calmodulin promoter. Integration into the *attB* genomic locus and subsequent sequencing was done using primers **P11 + P12**.

**Table 1.**
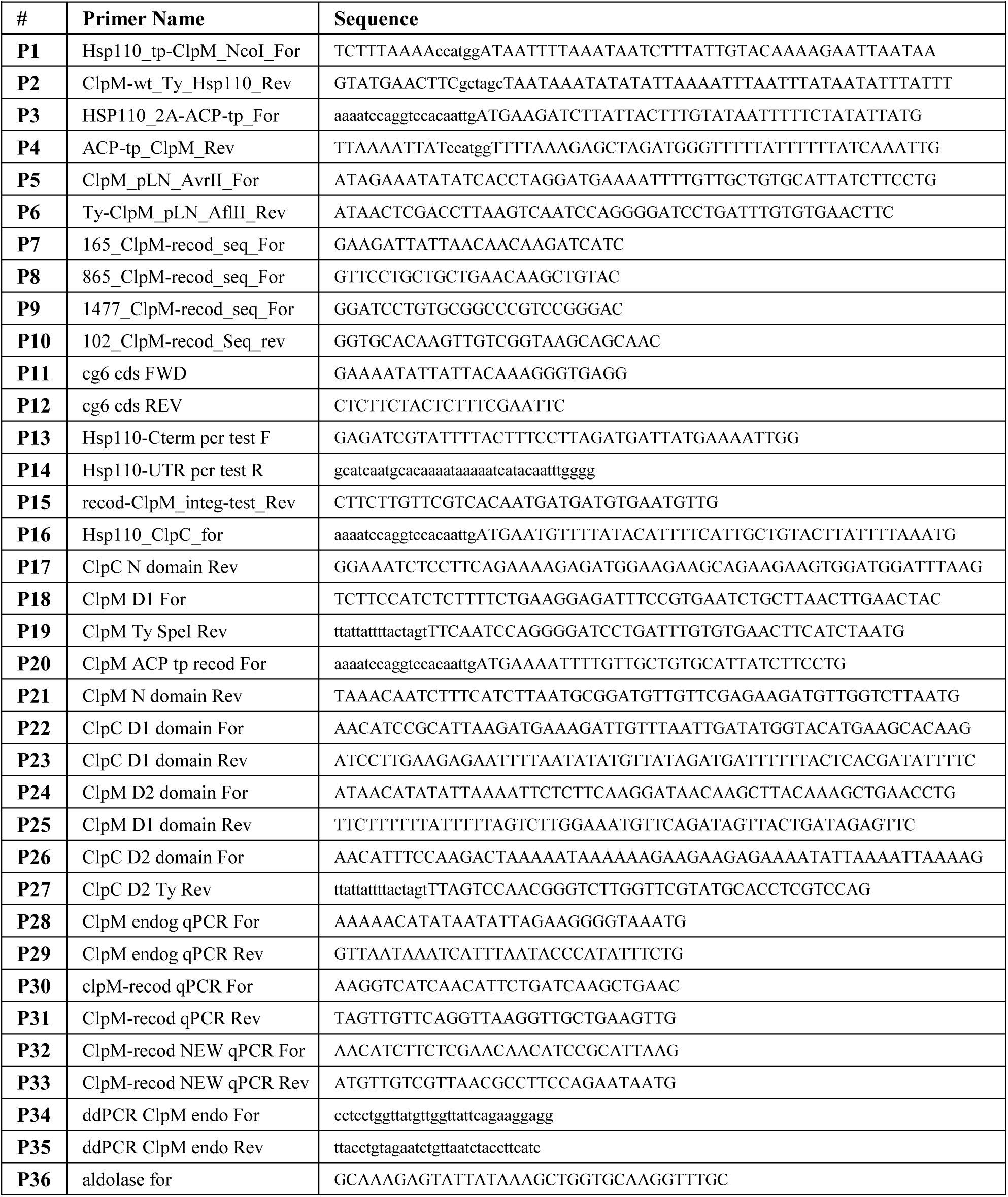
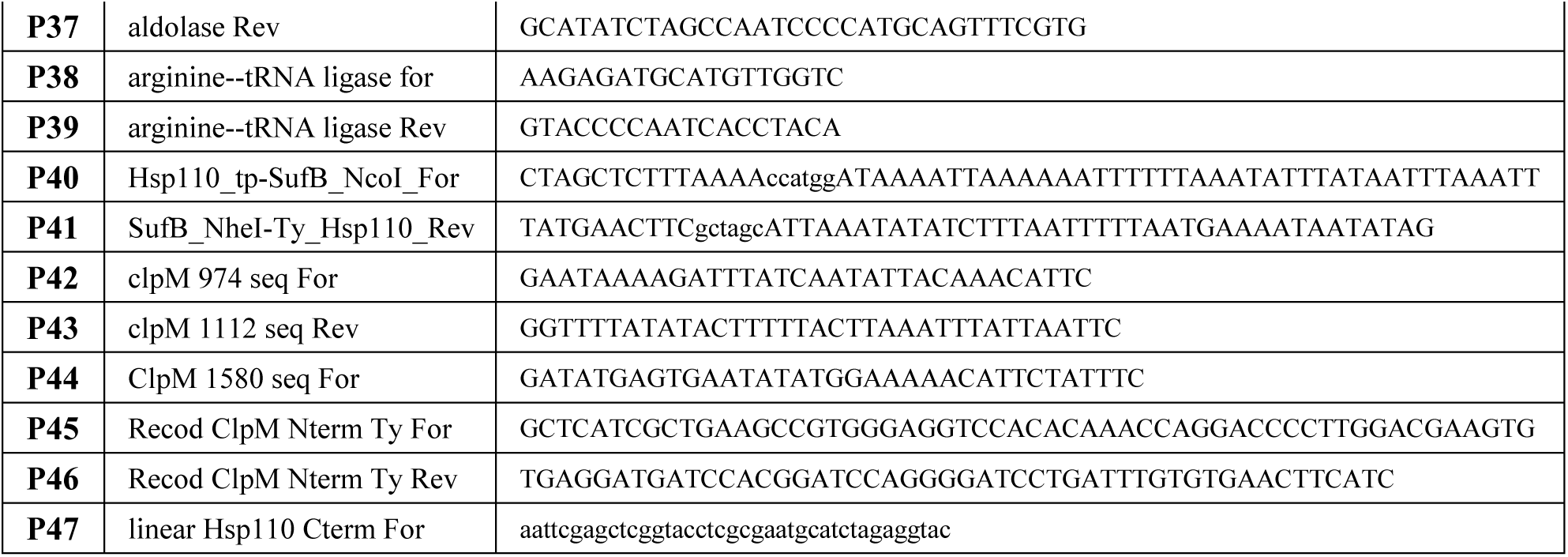
Primers used in this study.

To generate the *hsp110:clpM* transgenic parasites, recodonized Full-length ClpM was amplified using primers **P20 + P19** and cloned into puc57-hsp110 plasmid^22^. This vector contained homology regions corresponding to the final 429 bp of the *Pfhsp110c* gene (PF3D7_0708800) excluding the stop codon and followed by a T2A skip peptide. The construct was Sanger-sequenced using primers **P7 + P8 +P9 + P10.**

To generate the *hsp110:clpM^2xTy^* transgenic parasites, the puc57-*hsp110:clpM* was cut between the transit peptide and the beginning of the recodonized ClpM gene, and an additional Ty tag was inserted before the ATG start codon using primers **P45 + P46**. The construct was Sanger-sequenced using primers **P46 + P47**.

To generate the *hsp110:clpM^N-clpC^* transgenic parasites, The N-terminal domain of the ClpC gene (PF3D7_1406600) was amplified from genomic DNA using primers **P16 + P17** and D1+D2 domains of the recodonized ClpM gene were amplified using primers **P18 + P19**. The two products were conjugated together using PCR-sewing, and inserted as one piece into puc57-*hsp110* plasmid. To generate the *hsp110:clpM^D1-clpC^*transgenic parasites, the transit peptide and the N-domain of ClpM were amplified using primers **P20 + P21**, the D1 domain of ClpC was amplified from genomic DNA using primers **P22 + P23**, and the D2 domain of the recodonized ClpM was amplified using primers **P24 + P19**. The three products were conjugated together using PCR-sewing, and inserted as one piece into puc57-*hsp110* plasmid.

To generate the *hsp110:clpM^D2-clpC^* transgenic parasites, the transit peptide, the N-domain and the D1 domain of ClpM were amplified using primers **P20 + P25**, and the D2 domain of ClpC were amplified primers **P26 + P27.** The two products were conjugated together using PCR-sewing, and inserted as one piece into puc57-*hsp110* plasmid.

To generate the *hsp110:clpM^endo^* transgenic parasites, the endogenous sequence of the *PfclpM* gene (PF3D7_API03600) was amplified from genomic DNA using primers **P1 + P2**, and fused to the transit peptide which was amplified using primer **P3 + P4**. The two products were conjugated together using PCR-sewing, and inserted as one piece into puc57-*hsp110* plasmid, before a triple Ty C-terminal tag. The construct was Sanger-sequenced using primers **P42 + P43 + P44.**

To generate the *hsp110:sufB^endo^* transgenic parasites, he endogenous sequence of the *PfsufB* gene (PF3D7_API04700) was amplified from genomic DNA using primers **P40 + P41**, and cloned into puc57-hsp110 between a transit peptide and a Ty1 tag.

All of the HSP110-based constructs, were co-transfected with PfHsp110c guide RNA expressing plasmid (pUF1-Cas9-Hsp110-guide-ref), and integration into the correct genomic locus was verified and sequenced using primers **P13 + P14 + P15.**

### Cell culture and transfections

Parasites were cultured in RPMI medium supplemented with Albumax I (Gibco) and transfected as described earlier^49–51^. Synchronization of cultures were achieved by intermittent sorbitol and Percoll treatments. To generate *attB-CAM:clpM* transgenic parasites, 30 μg of the pLN plasmid were co-transfected with 15 μg pINT plasmid expressing the Bxb1 integrase for site-specific integration into the attB locus of (*wildtype_NF54^attB^*) parasites^37^. Selection was applied 48 h post-transfection using 2.5 μg/mL blasticidin (Sigma). Clonal lines were obtained by limiting dilution, and clone A4 was used for all subsequent experiments. To generate HSP110-based constructs (*hsp110:clpM, hsp110:clpM^x2Ty^, hsp110:clpM^N-clpC^, hsp110:clpM^D1-clpC^, hsp110:clpM^D2-clpC^, hsp110:clpM^endo^, hsp110:sufB)* a mix of two plasmids was transfected into *NF54 wildtype* parasites. For each different transfection the primers mix contained 2 plasmids; 30 μg of pUF1-Cas9-PfHsp110c-guide and 40 μg of the relevant marker-less puc57-Hsp110 repair plasmid with the corresponding transgene. Parasites’ transformation was carried out as described before^52^. Briefly, 400 μl packed erythrocytes were electroporated (0.32 kV, 925 μF; Gene Pulser II, Bio-Rad) together with a mixture of of purified plasmid DNA resuspended in 380 μl of CytoMix solution (120 mM KCl, 0.15 M CaCl2, 2 mM EGTA, 5 mM MgCl2, 10 mM K2HPO4/KH2PO4, and 25 mM HEPES [pH 7.6]). Schizont-stage parasites were added, and selection was applied 48 hours post transfection, using 1 μM DSM1 (BEI Resources), selecting only for Cas9 expression. DSM1 was removed from the culturing media once parasites clones were isolated using limiting dilutions. Clones were screened, and the clone exhibiting the most robust expression was selected for downstream experiments. Representative clones used were D4 (*hsp110:clpM*), E1 (*hsp110:clpM^D1-clpC^*), and H7 (*hsp110:clpM^D2-clpC^*). Generation of the *hsp110:clpC* and *hsp110:GFP* transgenes was described earlier^22^.

For stabilizing lower temperatures, parasites were incubated at 30°C, 32°C, or 34°C for 48 h in separate experiments, each performed in parallel with control cultures maintained at 37°C. To test for ER-associated degradation, parasites were treated with the proteasome inhibitor MG-132 (SigmaAldrich) at a final concentration of 200 nM for 6 h. For apicoplast ablation, parasites were treated with 40 μM Chloramphenicol (Goldbio) supplemented with 200 μM IPP (Isoprenoids LC).

### Western blot

Western blots were performed as described previously^22^. Briefly, parasites were collected and host red blood cells were permeabilized selectively by treatment with ice-cold 0.04% saponin in PBS for 15 min, followed by a wash in ice-cold PBS. Cells were lysed using RIPA buffer, sonicated, and cleared by centrifugation at 4°C. The antibodies used in this study were mouse anti-Ty, BB2 (inhouse purification), polyclonal rabbit anti-GFP, A-11122 (Invitrogen, 1:1000) and rabbit anti-aldolase ab207494 (Abcam, 1:3000). The secondary antibodies that were used are IRDye 680CW goat anti-rabbit IgG and IRDye 800CW goat anti-mouse IgG (LICOR Biosciences, 1:20,000). The Western blot images and quantifications were processed and analyzed using the Odyssey infrared imaging system software (LICOR Biosciences). Three biological replicates were used for quantification of protein expression levels using Prism 10.5.0 (GraphPad Software, Inc.).

### Growth assays and flow cytometry

Asynchronous parasite cultures were seeded at 0.5% parasitemia, and measured every 24 h by flow cytometry over 11 days. Cumulative parasitemia values were back-calculated by multiplying measured parasitemia by the relevant dilution factors. Parasitemia of the *wildtype* strain at the end of each experiment was set as the highest relative parasitemia and was used to normalize parasites growth. For flow cytometric analysis, 5 μL aliquots of culture were stained with 8 μM Hoechst 33342 (Thermo Fisher Scientific) in PBS. Fluorescence was measured by flow cytometry (CytoFlex S). Data were analyzed using Prism 10.5.0 (GraphPad Software, Inc), and growth curves were fitted to an exponential model. To measure multiplication rate, parasitemia values were log₁₀-transformed and plotted against time (days) to model exponential growth. The exponential phase was identified, and data points within this interval were fitted using linear regression in GraphPad Prism. The slope of the fitted line (m, day⁻¹) was taken as the parasite growth rate. The parasite multiplication rate over 48 hours (PMR₄₈) was calculated for each biological replicate as previously described^53^; using the equation PMR₄₈ = 10^(2m), corresponding to one full asexual replication cycle. PMR₄₈ values were calculated independently for each replicate and are presented as mean ± SEM. All experiments are representative of at least three independent biological replicates.

### Half-maximal effective concentration (EC50)

To generate an EC50 curve for chloramphenicol, asynchronous cultures of *wildtype*, *hsp110:clpM^D1-clpC^*, and *hsp110:clpM^D2-clpC^*were seeded at 0.5% parasitemia in a 96-well plate. Parasites were treated with chloramphenicol at concentrations ranging from 100 - 0.1 μM using serial dilutions. Parasitemia was measured after 120 h by flow cytometry using Hoechst 33342 staining. Dose–response curves were fitted using Prism (GraphPad Software, Inc.).

### Microscopy and image processing

For IFA, cells were fixed using a mix of 4% paraformaldehyde and 0.015% glutaraldehyde and permeabilized using 0.1% Triton-X100. Primary antibodies used are mouse anti-Ty1, BB2 (Invitrogen, 1:100), or rabbit anti-aldolase ab207494 (Abcam, 1:3000). Secondary antibodies used are Alexa Fluor 488 or Alexa Fluor 546 (Life Technologies, 1:100). Cells were mounted on Fluoroshield with DAPI (Sigma). The imaging was performed using a Nikon Spinning Disk confocal fluorescence microscope equipped with a 100x/1.4NA objective. Images were collected as Z-stack, and displayed as maximum intensity projection. Image processing, analysis and display were performed using Nis Elements Software (Nikon). Adjustments of brightness and contrast were made for display purposes. Blood smears were imaged using an upright Eclipse E200 Microscope (Nikon), equipped with x100 oil objective Type NVH, and captured using a digital Sight 1000 microscope camera and NIS-Elements Software (Nikon).

### RNA extraction, quantitative real time PCR (qRT-PCR), and Digital droplet PCR (ddPCR)

Total RNA was extracted using NucleoSpin RNA Prep Kit (Macherey-Nagel). RNA concentration and purity were assessed using NanoDrop spectrophotometer (Thermo Fisher Scientific). After normalizing the concentrations of all the samples, 15 µL of the total RNA (from a 40 µL eluate) were used to synthesize complementary DNA (cDNA) using qScript cDNA Synthesis Kit (QuantaBio). For both qRT-PCR and ddPCR, gene-specific primers were used to amplify the endogenous *PfclpM* gene (PF3D7_API03600, **P28 + P29, P34 + P35**), recodonized transgenic clpM gene (**P30 + P31, P32 + P33**), and the housekeeping genes: *aldolase* (PF3D7_1444800, **P36 + P37**) and *arginine-tRNA ligase* (PF3D7_1218600, **P38 + P39**) for normalization.

qRT-PCR reactions were prepared using the Myra automated liquid handling system in a final reaction volume of 20 μl, containing 10 μl of Luna Universal qPCR Master Mix, 1 μl of RNA sample, 0.5 μl each of forward and reverse primers (10 μM), and 8 μl of ultrapure water. Reactions were conducted using a Quant Studio real-time PCR system (Thermo Fisher Scientific). Each reaction was performed in triplicate. Relative expression levels were calculated using the ΔΔCt method, normalizing to the geometric mean of the housekeeping genes. Statistical analyses were performed using Prism 10.5.0. (GraphPad Software, Inc.)

Digital droplet PCR was performed using QX200 ddPCR EvaGreen Master Mix (Bio-Rad) with identical primer sets across all samples. Each reaction was performed once per biological replicate, and transcript levels were normalized to housekeeping genes. Statistical analyses were conducted using GraphPad Prism version 10.5.0 (GraphPad Software, Inc.). All experiments were repeated across three independent biological replicates.

### Bioinformatic analysis

To identify clpM homologs across plastid-containing organisms, a total of 15,231 complete plastid organelle genome records were retrieved from the NCBI Organelle Genome Resources database^54^ (https://www.ncbi.nlm.nih.gov/genome/organelle/) and classified into taxonomic groups using the ETE3 toolkit^55^ with NCBI taxonomy. Protein sequences were extracted from the CDS features of each genome using Biopython^56^. BLASTp^57,58^ was used to search for homologs using PfClpM (PF3D7_API03600.1, 767 aa) as query (E-value ≤ 1e-5). Borderline hits were manually inspected and excluded if they lacked significant similarity beyond the E-value threshold.

Sixteen representative organisms were selected to span key plastid-containing lineages (Apicomplexa, Chromerida, Stramenopiles, Rhodophyta, and Viridiplantae), with phylogenetic relationships retrieved from the Open Tree of Life synthetic tree^59^ using the OpenTree Python package^60^. The completeness of protein-coding gene annotation was assessed for all 16 organisms using BUSCO v6.0.0^61^ in protein mode, with the most specific available OrthoDB v12 lineage dataset for each organism. All proteomes scored ≥80% complete, except for Theileria parva (69.4%). Nuclear proteomes were obtained from UniProt Reference Proteomes^62^, with the exception of Chromera velia that was retrieved from VEuPathDB^63^. Plastid organelle genome sequences were retrieved from the NCBI Organelle Genome Resources database described above. To characterize the nuclear-encoded Clp chaperone repertoire across the 16 representative organisms, BLASTp analysis against nuclear proteomes was performed with PfClpC (PF3D7_1406600.1, 1341 aa) and PfClpM as queries (E-value ≤ 1e-5).

Protein domain architecture was analyzed using HMMER 3.4^64^ against the Pfam-A database^65^, retaining hits with i-Evalue ≤ 1e-5. To verify the functional integrity of each AAA ATPase domain, proteins were additionally searched for the canonical nucleotide-binding signatures of AAA+ ATPase domains: Walker A (GxxxxGK[S/T]) and Walker B (hhhhD[D/E]) motifs^66,67^. Nuclear-encoded proteins were retained only if they contained two functionally validated AAA ATPase domains. This analysis was performed for both nuclear and organelle-encoded proteins, and the resulting domain sequences were used for phylogenetic analysis.

For both phylogenetic trees, sequences were aligned using MAFFT v7^68^, trimmed using trimAl v1.4^69^, and maximum likelihood reconstruction was performed using IQ-TREE2 v2.3.6^70^ with ModelFinder for best-fit model selection and 300 nonparametric bootstrap replicates. Bayesian inference was additionally performed using MrBayes^71^. For the nuclear-encoded Clp chaperone tree (Fig. 1D), the best-fit model was LG+R5. For the AAA ATPase domain tree (Fig.6A),domain sequences were first extracted: for proteins with two AAA ATPase domains, D1 was defined from its start to the residue preceding D2, and D2 from its start to the C-terminus. For single-domain proteins, the region spanned from the domain start to the C-terminus. The best-fit model was LG+I+G4. All trees were visualised using iTOL^72^.

Pairwise sequence alignment analysis was performed between PfClpM and PfClpC using EMBOSS^73^, local alignment of the full-length proteins was performed using EMBOSS Water. Domain-level global alignments were performed using EMBOSS Needle for the D1 AAA ATPase domain (PfClpM 153-439 aa; PfClpC 506-913 aa) and the D2 AAA ATPase domain (PfClpM 440-767 aa; PfClpC 914-1341 aa), with domain boundaries as previously determined^21^. Alignments were visualized using ESPript 3.2^74^.

## Data Availability

All of the data described are included in the manuscript and Figures.

## Acknowledgements

We thank Boris Striepen for the BB2 hybridoma cell line; Josh Beck for pLN and pINT plasmids; Yuval Tabach for consultation on bioinformatic work. Florentin lab members, Oliver Caspari and Yiska Weisblum for critical reading and comments on the manuscript; VEuPathDB and PlasmoDB for scientific and community support. This work was supported by grants from the Israel Science Foundation 400/22 and 2786/22 to AF and HUJI start-up funds to AF. AQ and SKG are supported by PhD scholarships from Science Training Encouraging Peace (STEP), and AQ is also supported by Carole and Andrew Harper Diversity PhD Program. AF is supported by the Abisch-Frenkel Faculty Development Lectureship. All authors acknowledge support by The Kuvin Center for the Study of Infectious and Tropical Diseases.

## Supplemental information

Sup Figures: Fig. S1, S2, S3, S4.

## Preprint Server

BioRxiv

---

**Figure S1.**
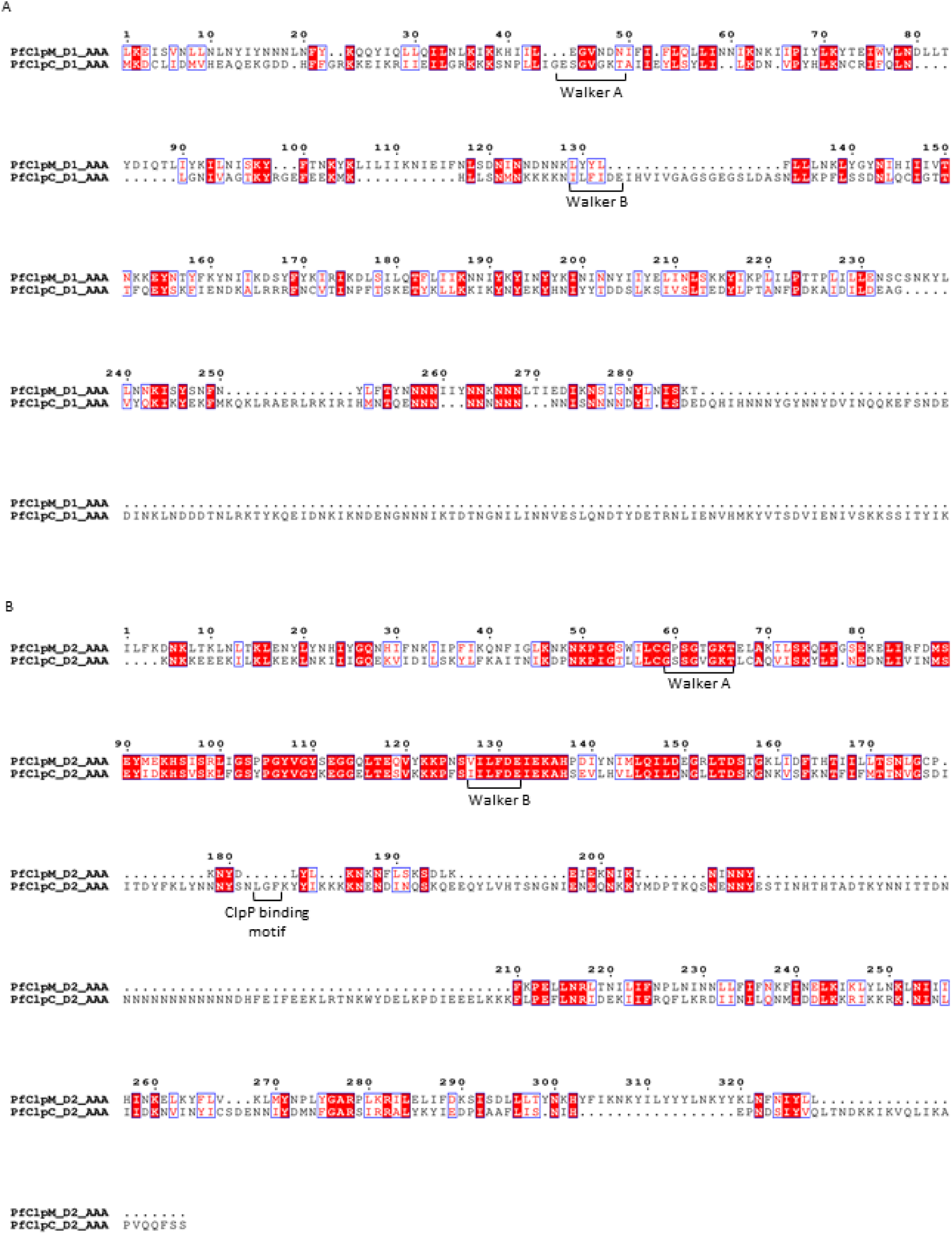
Pairwise sequence alignment of the D1 and D2 AAA ATPase domains of *P. falciparum* ClpM and ClpC. **A**. Alignment of the D1 AAA domain of *PfClpM* (aa 153-439) and *PfClpC* (aa 506-913); 15.3% identity, 30.1% similarity. **B**. Alignment of the D2 AAA domain of *PfClpM* (aa 440-767) and *PfClpC* (aa 914-1341); 28.8% identity, 43.6% similarity. Alignments were performed using EMBOSS Needle (Needleman–Wunsch algorithm; BLOSUM62, gap open 10, gap extend 0.5) and rendered with ESPript 3. Identical residues are shown on a red background, and similar residues are shown in red. Walker A (GxxxxGK[S/T]), Walker B ([VILMF]_4_DE) and ClpP-binding motifs are indicated.

**Fig. S2.**
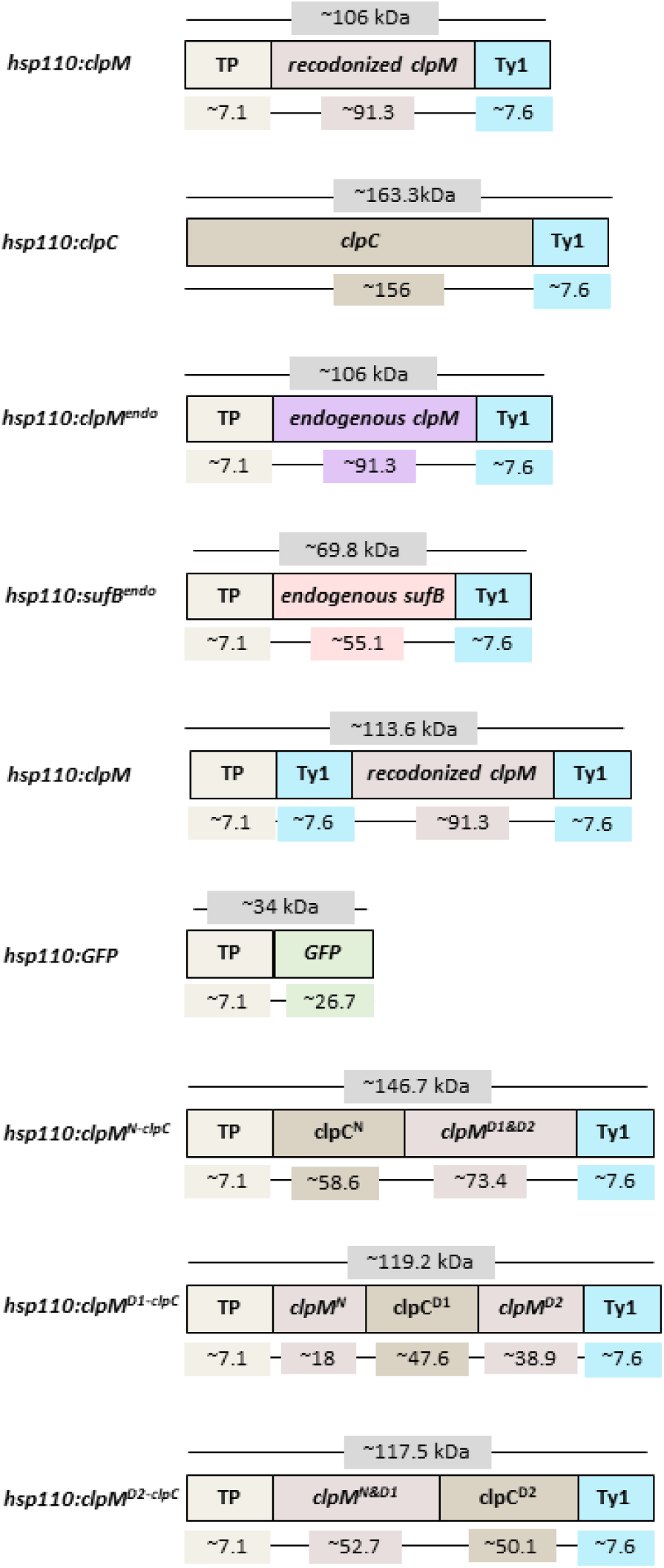
Predicted molecular weights of transgenic constructs generated in this study. Schematic summary of the predicted molecular weights of proteins expressed from the indicated transgenic parasite lines, including *hsp110:clpC*, *hsp110:clpM*, *hsp110:clpM^endo^*, *hsp110:sufB^endo^*, *hsp110:GFP*, *hsp110:clpM^N-clpC^*, *hsp110:clpM^D1-clpC^*, and *hsp110:clpM^D2-clpC^*. Predicted sizes were calculated based on the combined molecular weights of all parts in each construct.

**Fig. S3.**
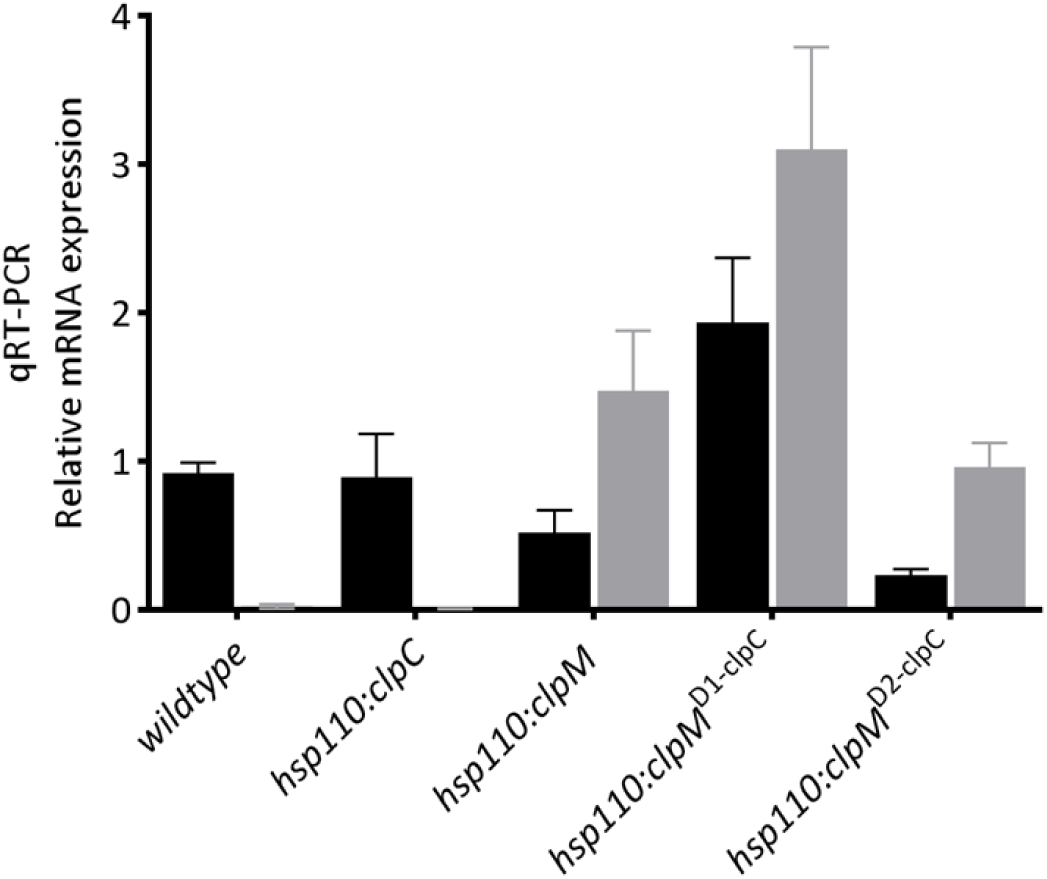
qPCR analysis performed on cDNA derived from saponin-isolated parasites. Expression was assessed using two primer sets: one detecting *endogenous pfclpM* (detected in all lines, including *wildtype* and *hsp110:clpC*), and a second specific to the recodonized constructs (detected in *hsp110:clpM, hsp110:clpM^D1-clpC^* and *hsp110:clpM^D2-clpC^* lines). Equal amounts of cDNA were used across samples. Error bars indicate standard deviation (SD) of two primers sets for the same gene. One representative experiment is shown, out of three biological replicates.

**Fig. S4.**
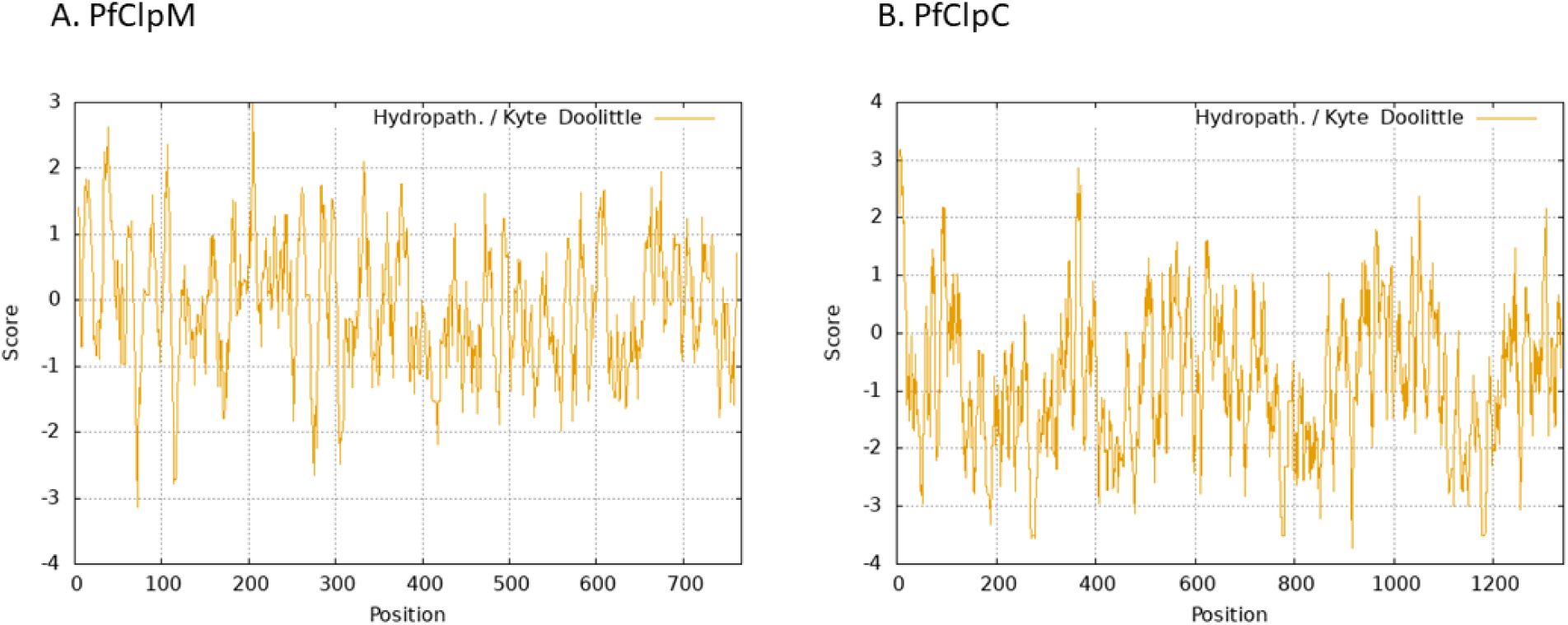
A Kyte-Doolittle hydrophobicity analysis of P. falciparum ClpM and ClpC. Amino acid sequences of PfClpM (A) and PfClpC (B) were analyzed for their hydrophobicity using ProtScale (an online tool: https://web.expasy.org/protscale/). Briefly, a sliding window (usually 5-25 residues) passes through the sequence, averaging the hydrophobicity to identify specific hydrophobic stretches, often used to predict transmembrane domains. Using this scale, the cut-off for hydrophobicity is typically above 4.5, suggesting that none is particularly hydrophobic.

